# RamanOmics Decodes Spatial Vibrational-Molecular Architecture and Rewiring in Aging and Repair

**DOI:** 10.64898/2025.12.04.692337

**Authors:** Ke Zhang, Xingjian Chen, Francesco Monticolo, Salvatore Sorrentino, Haochun Huang, Claire Callahan, Yueqing Qiao, Judy Zhou, Sonia Brodowska, Styliani Sapantzi, Jianhuan Qi, Yinghan Wu, Thai Nam Son Dang, Francesca Viggiani, Chia-Kang Ho, Yanxin Xu, Koseki J. Kobayashi-Kirschvink, Thang Mung, Hemali Phatnani, Zhixun Dou, Jeon Woong Kang, Peter T. C. So, Jian Shu

## Abstract

Aging and tissue repair involve multilayered and spatially heterogeneous remodeling across transcriptional, biochemical, and cellular dimensions, yet prevailing definitions rely on isolated molecular markers that obscure how biochemical and transcriptional states co-evolve in tissues. Here we present RamanOmics, a multimodal framework that integrates single-nucleus RNA sequencing (snRNA-seq), spatial transcriptomics, and label-free Raman imaging to map the spatial vibrational–biochemical and molecular architecture of aging and senescence directly in intact tissues. Applied to mouse lung and skin, RamanOmics generates spatially resolved biochemical–molecular maps revealing tissue-specific programs: lung senescent cells are enriched for extracellular matrix (ECM) remodeling and TGF-β signaling (*Serpine1, Dab2, Igfbp7*), whereas skin senescence is dominated by keratinization and barrier homeostasis modules (*Krt10, Lor, Sbsn*). Across tissues, we identify a conserved branched-chain fatty-acid-linked biochemical profile and Raman signature (1131-1135 cm⁻¹) that robustly marks *p21*⁺ senescent cells. To unify these layers, we develop a machine learning derived “multimodal barcode” that quantitatively integrates biochemical and transcriptional features, enabling non-destructive identification of senescence *in situ*. In a wound-healing model, RamanOmics further reveals coordinated reactivation of barrier-repair programs in senescent cells, marked by upregulation of *Krt10*, *Lor*, *Sbsn*, *Sfn*, and *Dmkn* together with matching increases in lipid-associated Raman signatures, confirming biological generalizability beyond steady-state aging. By directly integrating gene programs to spatial vibrational–biochemical states, RamanOmics provides a general framework and resource for scalable, multimodal profiling of cellular states.

## Main

Aging and tissue repair arise from coordinated multilayered regulation in gene expression, biochemical reactions, cellular state and tissue architecture, yet current frameworks reveal only a subset of this complex regulation. Single-cell transcriptomics and spatial genomics have transformed our understanding of aging, senescence, and regeneration by defining transcriptional profiles and programs associated with senescence, inflammation and repair^1,2^. However, transcription alone only reflects partial view of the multilayered regulatory networks and does not fully uncover the biochemical and metabolic composition of cells, such as lipid distribution, protein conformation, metabolite states and ECM chemistry, that together ultimately governs cell and tissue function^3^. This limitation creates a major gap in our ability to understand how the coordinated regulatory networks manifest within intact tissues.

Biochemical and vibrational imaging provide a unique view into the chemical-bond level architecture of cells, down to cellular or subcellular spatial resolution, capturing features such as C=O stretch, C–H, C–C bonds, lipid chain branching, protein secondary structure and nucleic-acid composition that are invisible to genomic technologies^4^. These vibrational modes, measured by Raman spectroscopy, report on the intrinsic physical and chemical properties of cells and their microenvironment while simultaneously collecting the full spectrum of chemical information optically, providing insights into both the composition and structure of molecules derived from the scattering of light as it interacts with molecular vibrations, and therefore complement molecular readouts in a fundamentally different way^4,5^. Changes in lipid saturation, protein folding or matrix chemistry often precede or occur independently of transcriptional changes, influencing membrane dynamics, signaling pathways and metabolic flux^6^. Thus, decoding biochemical and vibrational states and their links and coordination with gene expression programs is essential for understanding how cellular phenotypes arise, how they evolve with aging and how they reorganize during repair.

Despite their importance, vibrational-biochemical features have rarely been integrated with genome-wide molecular profiling *in situ*, in part because label-free biochemical imaging and high-dimensional molecular and genomic assays have historically operated in isolated experimental fields with limited crosstalk. Recently, we have pioneered and pushed the boundaries between these two domains and demonstrated that gene expression programs measured by single-cell RNA-seq and biochemical profiles measured by Raman imaging are intrinsically linked and associated and single-cell gene expression profiles can be predicted from label-free Raman images in live cells^7^. However, it has remained unclear how biochemical states relate to transcriptional programs across tissues, whether biochemical signatures of senescence are conserved across organs and how these states are dynamically rewired during repair and regeneration. Addressing these questions requires a framework that unifies biochemical, vibrational and molecular information at cellular resolution within intact tissues.

Here we develop RamanOmics, a multimodal platform that integrates snRNA-seq, STARmap *in situ* sequencing (STARmap-ISS), label-free hyperspectral Raman imaging, and machine learning to jointly map vibrational–biochemical and molecular architecture of aging and wound healing in mouse lung and skin. This multimodal analysis reveals coordinated lipid remodeling, ECM dysregulation, keratinization, and impaired DNA-repair programs as in-tissue hallmarks of senescence. With these integrated modalities, we develop a machine learning derived “multimodal barcode” that quantitatively fuses vibrational–biochemical and molecular features of senescence, reconstructing multimodal view of senescent cells. By directly coupling gene programs to the biochemical and vibrational states they encode, RamanOmics provides a conceptual and technological framework for dissecting the biochemical and molecular basis of aging and repair, and positions label-free multimodal tissue mapping as a promising strategy for non-destructive identification of senescence and cellular states *in vivo*.

## Results

### Multimodal characterization and identification of aging and senescence through RamanOmics

To investigate the cellular and molecular characteristics of aging in mouse tissues, we focused on lung and skin, which are key organs that function as protective barriers and are constantly and directly exposed to environmental factors (e.g., UV and air pollutants) contributing to senescence with age. We collected samples from 2- and 26-month-old mice. Nuclei were isolated from frozen OCT-embedded tissues for snRNA-seq from mouse lung (old, n= 3; young, n= 3) and mouse skin (old, n= 3; young, n= 3). Tissue sections from the same blocks were used for Raman imaging followed by spatial transcriptomics (old, n= 3; young, n= 3) (**Fig. 1a**). Raman imaging captured the subcellular full-spectrum of biochemical phenotypes, followed by imaging-based single-cell spatial transcriptomics (STARmap-ISS)^8^ to map cell states using a curated panel of 890 genes including 700 cell type marker genes and 190 senescence and aging-specific markers, and finally we employed *in situ* hybridization (STARmap-ISH) to validate newly identified senescent markers (**Table S1**). We successfully integrated these multimodal profiles using our newly developed platform, RamanOmics, and identified multimodal signatures of aging and senescence, which enabled the accurate multimodal spatial identification and characterization of vibrational-biochemical and molecular architecture of cellular senescence across tissues and ages at single-cell resolution (**Fig. 1a**).

**Figure 1.**
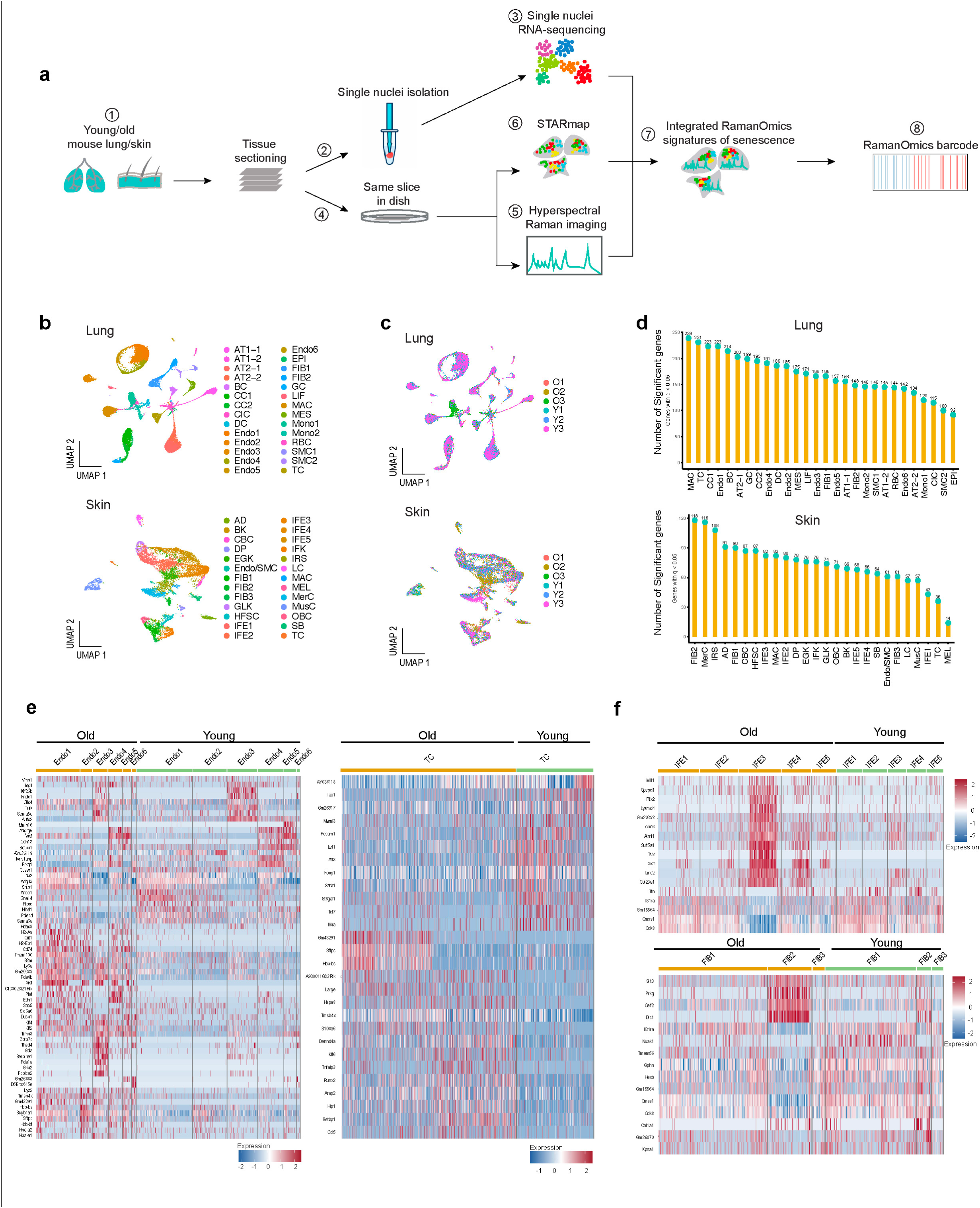
RamanOmics workflow and dynamic of gene regulation across ages and tissues. **a**, Detailed workflow of the RamanOmics: Young (2 months) and old (26 months) mouse lung and skin were collected from natural old mice and embedded into OCT (1); Adjacent sample slices from OCT were used for snRNA-seq (2) or fixed in Quartz bottom dishes (4); 10X single-nuclei transcriptome sequencing and RNA-seq analysis was performed (lung: old, n= 3; young, n= 3; skin: old, n= 3; young, n= 3) (3); Fingerprint of Raman spectra (600-1800 cm⁻¹) captured by Raman microscopy (lung: old, n= 3; young, n= 3; skin: old, n= 3; young, n= 3) (5); The same sample slices captured by Raman microscopy then proceeded to STARmap imaging to acquire the spatial transcriptomic information (6); Integrated RamanOmics signatures of senescence were generated by integrating snRNA-seq and Raman spectra information, leveraging STARmap spatial transcriptomic information (7). RamanOmics barcodes were developed by extracting important features from Integrated RamanOmics signatures of senescence to characterize and visualize the senescent or non-senescent cells intuitively (8). **b**, UMAP plots of snRNA-seq based on clustering. Cells are colored according to cluster identification. Cluster composition can be found in Supplementary Table 2. **c**, UMAP plots showing all sequenced single-nuclei samples from mouse lung and skin. Cells are colored according to samples. **d**, Lollipop plots showing the number of significantly altered genes discovered by EMD analysis (false-discovery rate (FDR)<0.05) per cell type in old versus young mouse tissues, ranked by the total number of significant features. **e**, Heatmaps displaying representative DEGs from snRNA-seq comparing old versus young mouse Endo (left panel) and TC (right panel) in the lung at sub-cell type level. The color key from blue to red indicates low to high gene expression levels. Endo, Endothelial cell; TC, T cell. **f**, Heatmaps displaying representative DEGs from snRNA-seq comparing old versus young mouse IFE (upper panel) and FIB (bottom panel) in the skin at sub-cell type level. The color key from blue to red indicates low to high gene expression levels. IFE, interfollicular epidermis; FIB, fibroblast cells.

### Distinct aging signatures across mouse lung and skin

To characterize the gene expression programs of aging and senescence, we generated high-quality single-nucleus transcriptomes from 35,474 mouse lung cells and 12,128 mouse skin cells, that passed stringent quality control (QC) criteria and filtering (**Extended data Fig. 1a**). Using principal component analysis (PCA)-based dimensionality reduction and subsequent clustering, we identified 28 major clusters in the lung samples and 26 in the skin samples by label transfer using Seurat, and manually curated cell type markers (**Fig. 1b, c, Extended data Fig. 1b, c**). We first analyzed transcriptomic alterations between young and old at the tissue level by performing gene sets enrichment analyses across all major cell types in mouse lung and skin. In the lung, aging was associated with widespread upregulation of genes involved in immune activation and inflammatory signaling, such as “leukocyte cell-cell adhesion” and “regulation of T cell activation”, across Alveolar type 1/2 cell (AT1/2), endothelial cell (Endo), and monocyte (Mono) (**Extended data Fig. 1d**). Concurrently, downregulated genes in the old lung were enriched in pathways related to transcriptional repression, epithelial proliferation, and ECM organization, particularly in T cells (TC), fibroblasts (FIB), and Endo (**Extended data Fig. 1e**). These results suggest that aging in the lung is characterized by chronic immune activation and downregulation of epithelial proliferative and ECM-organization pathways, consistent with reduced regenerative capacity and compromised structural maintenance.

In old mouse skin, upregulated genes were notably enriched in “ribonucleoprotein complex binding”, suggesting increased activity in RNA processing and protein synthesis machinery (**Extended data Fig. 1f**). In contrast, downregulated genes showed strong enrichment in pathways related to “response to metal ion” and “muscle contraction” across several cell types including Inner Root Sheath cell (IRS), Interfollicular Epidermis (IFE) cell, Interfollicular Keratinocytes (IFK), Muscle cell (MusC) (**Extended data Fig. 1g**). These changes reflect impaired ion homeostasis and muscular function, highlighting age-related declines in structural and metabolic integrity.

To further quantify transcriptional divergence with aging, we applied Earth Mover’s Distance (EMD), which measures the ‘moving cost’ between two distributions, to compare young and old expression distributions for each gene within each cell type^9^. In the lung, Endo and TC exhibited pronounced aging-associated transcriptomic remodeling, whereas in the skin, FIB and IFE cells showed substantial shifts (**Fig. 1d**). These population-level changes were accompanied by reduced cell abundance in Alveolar type 2 subtype 2 (AT2-2) and Endo2/3 cells, and a concurrent increase in cell numbers in TC and monocyte subtype 1 (Mono1) in the old lung (**Extended data Fig. 1h**). Similarly, old skin showed a decline in cell abundance in Merkel cells (MerC) and outer bulge cells (OBC), with increased proportions of TC, Langerhans cells (LC), adipocytes (AD), and early granular keratinocytes (EGK) (**Extended data Fig. 1h**).

Together, these results reveal tissue-specific aging patterns: the lung shows immune activation and vascular remodeling with diminished epithelial renewal, while the skin exhibits metabolic decline and impaired ion homeostasis alongside partial preservation of epithelial programs.

### Cell type specific transcriptional programs in aging

At the cell type level, we focused on those cell types that exhibited significant aging-associated transcriptional shifts in the EMD analysis. In old lung endothelial cells, we observed upregulation of gene modules involved in transcriptional regulation (*Klf2*, *Klf4*)^10^, antigen processing and presentation (*H2-Aa, H2-Eb1, Cd74, Ciita*)^11,12^, and ECM remodeling (*Timp3*, *Serpine1*)^13,14^ (**Fig. 1e**, **left**). Old T cells showed increased expression of inflammatory signaling and immune stress (*Tnfαip3*, *Ccl5*)^15,16^, and decreased expression of transcription factors associated with T cell activation (*Foxp1*, *Lef1*, *Aff3*)^17,18^ (**Fig. 1e**, **right**). In skin fibroblasts (FIB), particular in FIB2, aging led to upregulation of fibrotic and structural remodeling pathways (*Slit3*, *Dlc1*, *Prkg1*)^19–21^ (**Fig. 1f, bottom**). IFE cells similarly exhibited upregulation of metabolic-(*Sult5a1*, *Gpcpd1*)^22,23^ and structural-related genes (*Col23a1*)^24^, reflecting widespread activation of stress adaptation and regenerative programs (**Fig. 1f, top**).

Together, these aging signatures delineate tissue- and cell type-specific trajectories of inflammatory, metabolic, and structural remodeling in old lung and skin, indicating that aging is driven by distinct programs across cell types rather than a uniform tissue-wide shift.

### Age-dependent functional reprogramming of senescence

We identify senescent cells utilizing the canonical senescence marker *p21*, which initiates senescence by inhibiting cell cycle progression through the *p53* pathway^25^. Importantly, it has been demonstrated that clearance of *p21^+^* cells can reverse senescence-associated phenotypes, a phenomenon not observed with *p16^+^* cells clearance, further underscoring the unique function of *p21* in this process^26^. We identified *p21*^+^ senescent cells enriched in specific cell types, including AT2, Endo, and macrophages (MAC) in the lung (**Fig. 2a**), and IFE, basal keratinocytes (BK), and inner root sheath (IRS) in the skin (**Fig. 2b**).

**Figure 2.**
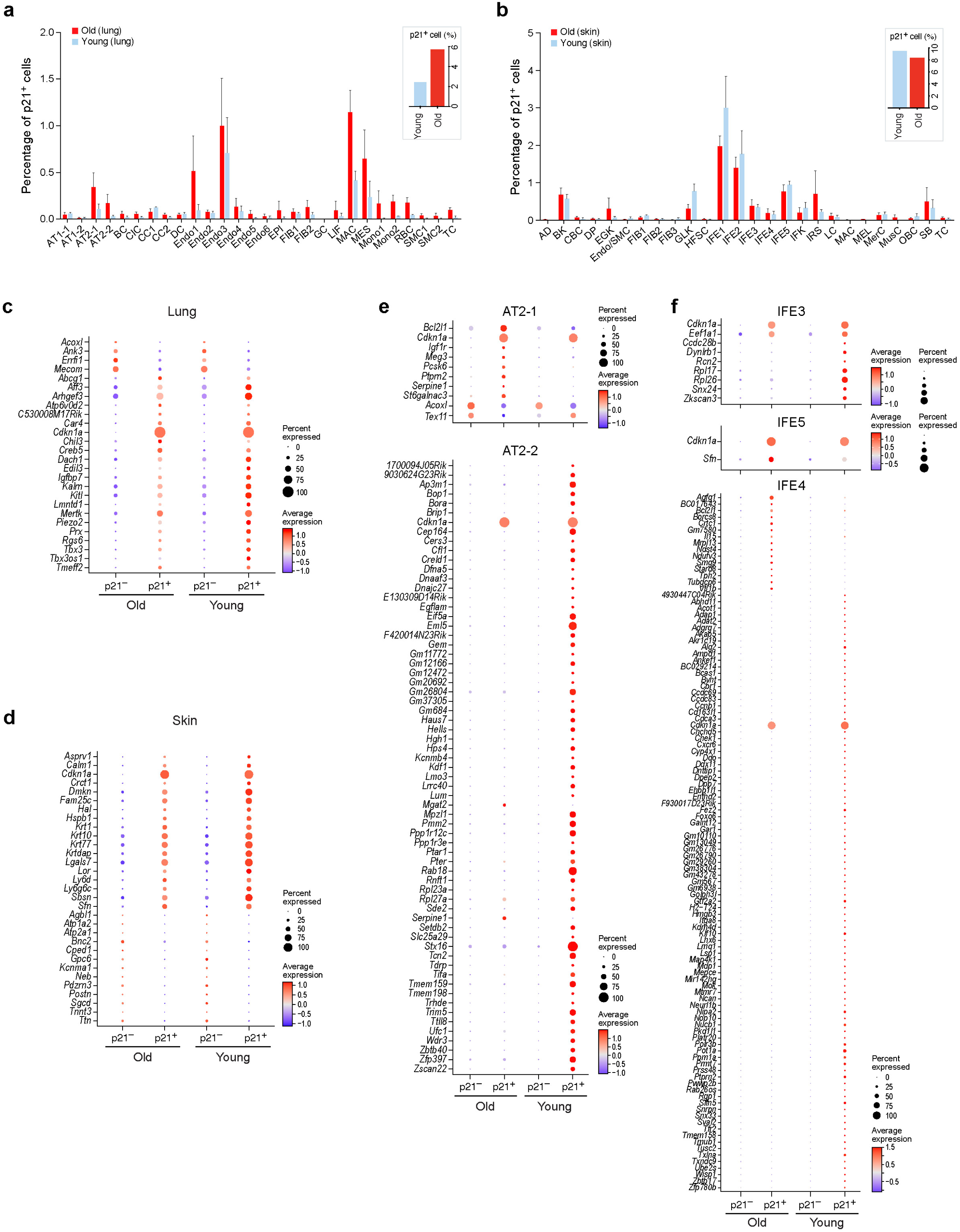
Cell type specific senescence programs revealed by differential expression profiling in aging lung and skin. **a-b**, Percentage of *p21*^+^ senescent cells at entire tissue and cell type resolution in mouse lung (**a**) and skin (**b**) from snRNA-seq results. The numerical results are reported in Supplementary Table 2. **c-d**, Dot plots showing shared DEGs between old and young samples in mouse lung (**c**) and skin (**d**). Average expression scores were calculated using log-normalized and scaled data (FDR<0.05). The numerical details are reported in Supplementary Table 6. **e-f**, Dot plots showing DEGs specifically enriched in AT2-1 and AT2-2 (**e**), IFE3, IFE4 and IFE5 (**f**) in old and young samples. Average expression scores were calculated using log-normalized and scaled data (FDR<0.05). The numerical details are reported in Supplementary Table 6.

In our comprehensive analysis of *p21*^+^ senescent cells from both young and old mouse lung tissues, we identified the shared up-regulated genes across ages including *Igfbp7*, *Abcg1*, *Prx*, *Tbx3* and *Mertk* (**Fig. 2c**), reflecting conserved programs related to oxidative stress, lipid metabolism, and apoptotic clearance^27–31^. In old lung, senescent cells upregulate *Serpine1*, *Dab2*, and *Mrc1* and downregulate *Acoxl*, *Aldh1a1*, and *Prdx6*, indicating a fibrosis- and inflammation-prone state^13,32,33^ with impaired lipid metabolism and antioxidant defense^34–36^ (**Extended data Fig. 2a, left**). In young lung, senescent cells upregulate *Dock10* and *Vav3* and downregulate surfactant/repair genes (*Sftpb*, *Sftpc*, *Evt5*), consistent with a more transient, reparative senescent program^37–41^ (**Extended data Fig. 2a, right**).

In mouse skin, senescent cells from both age groups commonly upregulate genes involved in epidermal barrier maintenance (*Krt1, Krt10, Lor, Sfn*) (**Fig. 2d**), indicating preserved epithelial defense^42–45^. Young skin specific senescent cells express differentiation and immune-interaction genes (*Lce1a2, Flg2, Skint5, Btc*), highlighting active repair and immune readiness^46–49^ (**Extended data Fig. 2b, right**). In contrast, old skin senescent cells show downregulation of lipid metabolism (*Cd36*, *Cidec*)^50,51^, and DNA repair genes (*Eepd1*)^52^ (**Extended data Fig. 2b, left**), reflecting diminished metabolic and regenerative function. These findings illustrate a shift from a responsive senescent state in young tissues to a chronic, dysfunctional one in old tissues.

### Cell type-specific programs of senescence across ages

Subsequently, we analyzed AT2 cells, which support gas exchange and repair by serving as progenitors for AT1 cells, particularly during lung injury or senescence^53,54^. In both AT2-1 and AT2-2 cells, old senescent cells showed consistent upregulation of *Serpine1* (**Fig. 2e**), a *p53* pathway-linked inhibitor of AT2 renewal and epithelial repair^55,56^. In AT2-1 of old sample, senescent cells also upregulated *Bcl2l1* and *Igf1r*, reflecting enhanced survival and apoptosis resistance^57^, while *Acoxl*, involved in fatty acid metabolism^36^, was downregulated across all senescent cells (**Fig. 2e**). The DNA repair gene *Tex11* was elevated only in young senescent AT2-1 cells, and young AT2-2 senescent cells similarly enriched DDR genes such as *Brip1*, *Bora*, *Hells*, and *Cep164* (**Fig. 2e**), collectively indicating age-related loss of DDR activity^58–62^.

In mouse skin, through the analysis of senescent IFE, which has the most abundant senescent cells in mouse skin (**Fig. 2b**). We identified senescent IFE5 and IFE3 showed enrichment of *Sfn* and *Eef1a1,* respectively (**Fig. 2f**), supporting ECM remodeling and protein synthesis^45,63^. In contrast, senescent IFE4 of young sample upregulated *Chek1*, *Ccnb1*, and *Cdca3*, indicating active DNA damage response and cell cycle regulation, reflecting a more dynamic and reparative senescent state^64–66^ (**Fig. 2f**).

Our findings reveal that senescent cell function shifts with age in a cell type- and tissue-specific manner: young tissues retain active DNA repair, while old tissues favor survival and inflammation, reflecting diminished repair capacity and metabolic flexibility.

### Increased cell-cell interactions and ECM remodeling in aging

Next, we examined how aging alters intercellular communication. We first assessed interactions of lung endothelial cells and T cells, and skin IFE cells and fibroblasts with other cell types, as these populations showed prominent age-related transcriptomic alterations (**Fig. 1d**). In old lung, endothelial cell subtypes demonstrated a global reduction in cell-cell interactions, including diminished endothelial self-communication (**Extended data Fig. 3a**). In contrast, we observed increased interactions between T cells and monocytes, dendritic cells, and AT2-2 cells (**Extended data Fig. 3a**). Similarly, in old skin, FIB and IFE cells showed decreased interactions with most other cell types yet exhibited enhanced communication between IFE and sebocytes (**Extended data Fig. 3b**).

To further explore how these changes translate into signaling dynamics, we analyzed age-associated pathway activity across lung and skin tissues. In the old lung, TGF-β signaling was prominently upregulated in Endo5, SMC1/2, and FIB populations, while AT1-1 and AT2-1 cells displayed elevated collagen signaling relative to young samples (**Extended data Fig. 3c**). These pathways are classically associated with fibrosis, vascular remodeling, and senescence, suggesting a shift toward a pro-fibrotic and inflammatory microenvironment with age. In the old skin, Notch signaling was enhanced in FIB3, IFE3 and IFK populations, whereas BMP signaling increased in OBC, FIB3 and IFE3 cells (**Extended data Fig. 3d**), indicating a transition toward differentiation, matrix remodeling, and altered epithelial homeostasis^67–70^.

To examine how these age-related signaling changes intersect with cellular senescence, we next focused on cell-cell interactions involving senescent cells. In the lung, *p21*^+^ AT2-2 and T cells exhibited increased interactions with both senescent and non-senescent cells compared to their younger counterparts (**Extended data Fig. 4a**), suggesting that senescent cells actively engage in shaping their local microenvironment. In contrast, in old skin, *p21*^+^ IFE4 and Langerhans cells predominantly increased interactions with non-senescent neighbors (**Extended data Fig. 4b**), possibly reflecting their roles in epithelial surveillance and immune crosstalk.

We further analyzed signaling pathway activation in senescent cells to understand the functional implications of these interactions. In old lung, senescent AT2-2 cells, T cells, macrophages, and monocytes showed heightened TGF-β signaling, while senescent AT2-2 cells, T cells, monocytes, and FIB2 displayed increased collagen signaling compared to young tissues (**Extended data Fig. 4c**). These pathways, known drivers of ECM remodeling, fibrosis, and senescence, point to a microenvironment that sustains and amplifies senescent cell states. Similarly, in the old skin, senescent IFE and IFK cells exhibited elevated Notch and BMP signaling (**Extended data Fig. 4d**), consistent with enhanced epithelial differentiation and remodeling during cutaneous aging^67,70^.

### Spatially resolved aging and senescent signatures across tissues and ages

To overcome the lack of spatial information of snRNA-seq, we employed STARmap-ISS to target 890 genes in space (lung: old, n= 3; young, n= 3; skin: old, n= 3: young, n= 3) (**Fig. 1a**; **Table S1**). Using label transfer from snRNA-seq references, we successfully identified 16 distinct cell types in the lung and 20 in the skin (**Fig. 3a, b, Extended data Fig. 5a, b**). To visualize spatial distribution and assess regional differences between young and old tissues, we generated spatial maps of the major cell types (**Extended data Fig. 5c, d**). The spatial distribution of these cell types faithfully reflected the native tissue architecture, demonstrating the accuracy and reliability of our single-cell spatial transcriptomic data.

**Figure 3.**
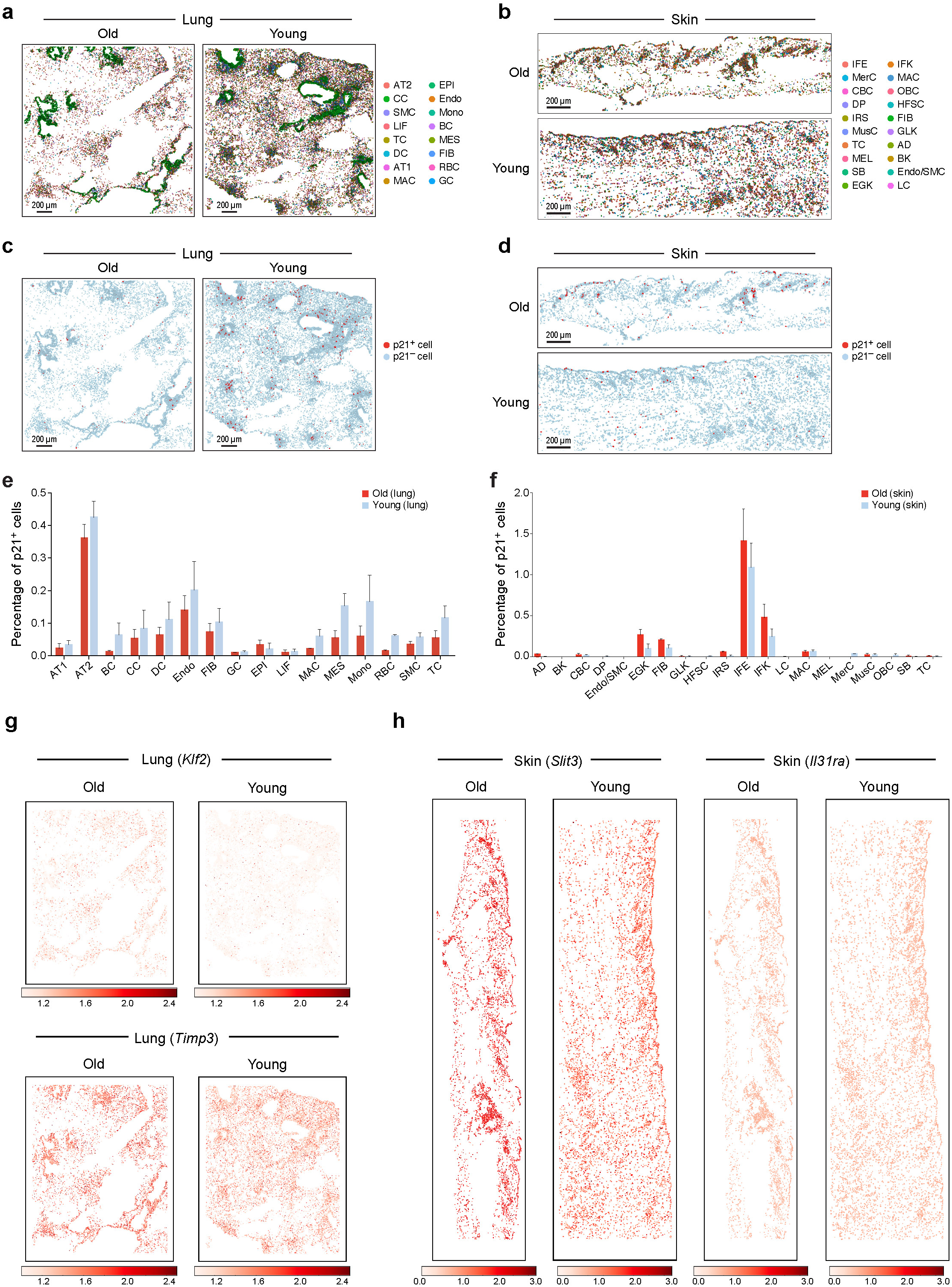
Spatial transcriptomic landscape of senescent cells in mouse lung and skin at different ages. **a-b**, snRNA-seq derived major cell types identified by STARmap-ISS (890 genes) via label transferring on old and young samples in mouse lung (**a**) and mouse skin (**b**). Cells are colored according to cell type identification. Cluster composition can be found in Supplementary Table 2. **c-d**, Spatial distribution of *p21*^+^ senescent cells in old and young samples in mouse lung (**c**) and mouse skin (**d**). **e-f**, Percentage of *p21^+^* senescent cells at cell type resolution in mouse lung (**e**) and skin (**f**) from STARmap-ISS result. The numerical results are reported in Supplementary Table 2. **g-h**, Spatial visualization of imputed expression of aging markers identified by snRNA-seq in mouse lung (**g**; *Klf2, Timp3*) and mouse skin (**h**; *Slit3, Il31ra*) across young and old sections. Values are expression-scaled per gene, with higher values indicating higher predicted expression

To assess senescent cell distribution across cell types, we calculated the percentage of *p21^+^* senescent cells in each annotated population. In mouse lung, AT2 cells exhibited the highest proportion of senescent cells (0.31%-0.51%), followed by endothelial cells (0.06%-0.36%), then immune cells such as dendritic cells (0.02%-0.10%), T cells (0.01%-0.17%) (**Fig. 3c, 3e; Table S2**). In mouse skin, the highest senescent cell composition was observed in IFE (0.53%-1.55%), followed by IFK (0.04%-0.64%) and FIB (0.07%-0.20%) (**Fig. 3d, f; Table S2**). Next, we mapped the spatial distribution of *p21*⁺ senescent cells across tissues and ages. Although we did not observe distinct spatial differences between senescent cells in young and old lung (**Fig. 3c**), we identified a higher concentration of senescent cells in the epidermal region of old skin (**Fig. 3d**). This finding is consistent with our earlier discovery that the IFE and IFK compartments have a higher proportion of senescent cells across different age groups (**Fig. 3f**). As the outermost layer of the skin, epidermal cells are particularly vulnerable to chronic UV exposure and oxidative stress, both of which are major contributors to skin senescence.

Next, we enhanced our spatial transcriptomic resolution from 890 genes to a whole-transcriptome scale by imputing single-cell transcriptomic profiles using Tangram^71^. As a result, we successfully imputed gene expression profiles for 19,812 genes across 46,932 cells in our old mouse lung and for 18,931 genes across 32,574 cells in our young mouse lung STARmap dataset. To assess imputation accuracy, we compared imputed and STARmap-ISS measured expression of *Scgb1a1* (club cell marker) and observed strong concordance in spatial distribution (**Extended data Fig. 6a**). For mouse skin, we imputed gene expression profiles for 19,294 genes in 13,153 cells from old skin and 18,464 genes in 18,583 cells from young skin. Similarly, imputed *Krt14* (canonical marker of basal keratinocytes) expression aligned well with observed distribution (**Extended data Fig. 6b)**, this result indicated robust concordance between the imputed and measured data and demonstrated the high accuracy and reliability of the imputation method.

Through our snRNA-seq analysis, we identified novel aging markers for old mouse lung (e.g., *Timp3*, *Klf2*) (**Fig. 1e**), and canonical senescent markers, *Igfbp7* and *Serpine1*, as well as potential novel senescent markers *Abcg1* and *Dab2* (**Fig. 2c, Extended data Fig. 2a**). We visualized the expression of these markers in space and found *Klf2* and *Timp3* were upregulated in old lung tissue, and the expression of *Timp3* was highly localized to airway-adjacent regions in old mouse lung tissue (**Fig. 3g**). Recent multi-organ proteomics profiling confirmed that *Timp3* expression increases with age across nine human tissues, including the lung, and was incorporated as a core component of the tissue-specific aging clocks^72^. For the senescent markers (e.g., *Igfbp7*, *Serpine1*, *Dab2*), there is no specific spatial distribution of these genes in mouse lung tissues (**Extended data Fig. 6c, e**), which is correlated with the *p21^+^* cells distribution in space. Together, *Igfbp7* and *Dab2* function as TGF-β–dependent mediators that reinforce senescence-associated transcriptional programs and couple *p21*-dependent cell-cycle arrest to aging-associated insulin resistance and ECM remodeling that drive tissue dysfunction^73–75^.

In mouse skin, we identified that *Slit3*, *Il31ra* were upregulated in old mouse FIB and IFE (**Fig. 1f, 3h**), while *Sbsn*, *Krt10, Lor,* and *Dmkn* were significantly upregulated in senescent cells from both young and old skin (**Fig. 2d**), but their distribution in space showed no significant difference between young and old, aligning with the snRNA-seq that they are enriched in senescent cells of both young and old skin tissues (**Extended data Fig. 6d, f**). These findings suggest that the upregulation of these genes in senescent skin cells shows promoted keratinocyte differentiation and epidermal barrier maintenance.

### Increased cell size in senescence

To assess how aging and senescence affect cellular morphology, given that size changes are key indicators of altered cellular function, metabolic state, and structural adaptations during aging across different tissues^76^, we analyzed cell sizes using single-cell segmentation from registered STARmap data. Overall, cells in old tissues were significant smaller than those in young tissues (lung: 8.84±3.86 vs 9.84±3.90, p-value (p)= 3.55E-64; skin: 11.93±6.51 vs 12.84±6.24, p= 6.58E-23) (**Extended data Fig. 7a; Table S3**); however, no significant differences in cell size were observed in Endo and IFE cells across age groups (**Extended data Fig. 7d**). In contrast, *p21^+^* senescent cells were consistently larger than their non-senescent counterparts in old skin tissues (**Extended data Fig. 7e**). In the lung, only *p21^+^* senescent mesenchymal cells (MES) showed a significant increase in size compared to non-senescent cells (13.6000±4.1593 vs. 9.2718±3.8256, p= 2.24E-02) (**Extended data Fig. 7b; Table S3**). In the skin, both global *p21^+^*senescent cells (13.8242±8.2484 vs. 11.8921±6.4650, p= 2.70E-02) and *p21^+^* senescent IFE cells (16.0952±9.9236 vs. 12.8356±6.9811, p= 3.04E-02), were significantly larger than their non-senescent counterparts in old tissues, and a similar trend was observed in young skin, where global *p21^+^* senescent cells (16.8182±8.4427 vs. 12.7565±6.1567, p= 1.76E-09) were markedly larger than non-senescent cells (**Extended data Fig. 7e; Table S3**). Interestingly, comparisons between senescent cells in young and old tissues revealed a size reduction in old skin tissues (13.8242±8.2484 vs. 16.8182±8.4427, p= 2.22E-03) (**Extended data Fig. 7c**). These findings suggest that, while senescent cells generally exhibit increased size in skin tissues, aging may drive a reduction in cell size, potentially indicating age-related morphological and functional changes. This size variation could reflect differential senescence dynamics across tissues and age groups, highlighting tissue-specific adaptations during aging.

### Biochemical landscape and barcode of senescence with label-free hyperspectral Raman imaging

Raman microscopy captures the biochemical fingerprint of cells and tissues without the need for extensive sample preparation or labeling, which preserves the integrity of the biological sample and comprehensively reports the vibrational energy levels of chemical bonds in molecules and biochemicals in a non-destructive and label-free manner with subcellular spatial resolution^77^. We conducted the hyperspectral Raman imaging (600-1800 cm⁻¹, 873 dimensions with a pixel size of 3 µm) for the same samples before we performed STARmap-ISS on them (**Fig. 1a**). To align these two different modalities, we performed registration using the DNA signal at 791 cm⁻¹ wavelength from Raman imaging, and DAPI staining from STARmap-ISS (**Fig. 4a**)^78^. After registration, we identified 15,201 and 11,069 overlapping cells from Raman imaging and STARmap-ISS of mouse lung and mouse skin, respectively (**Table S2**), demonstrating appropriate alignment between these two modalities.

**Figure 4.**
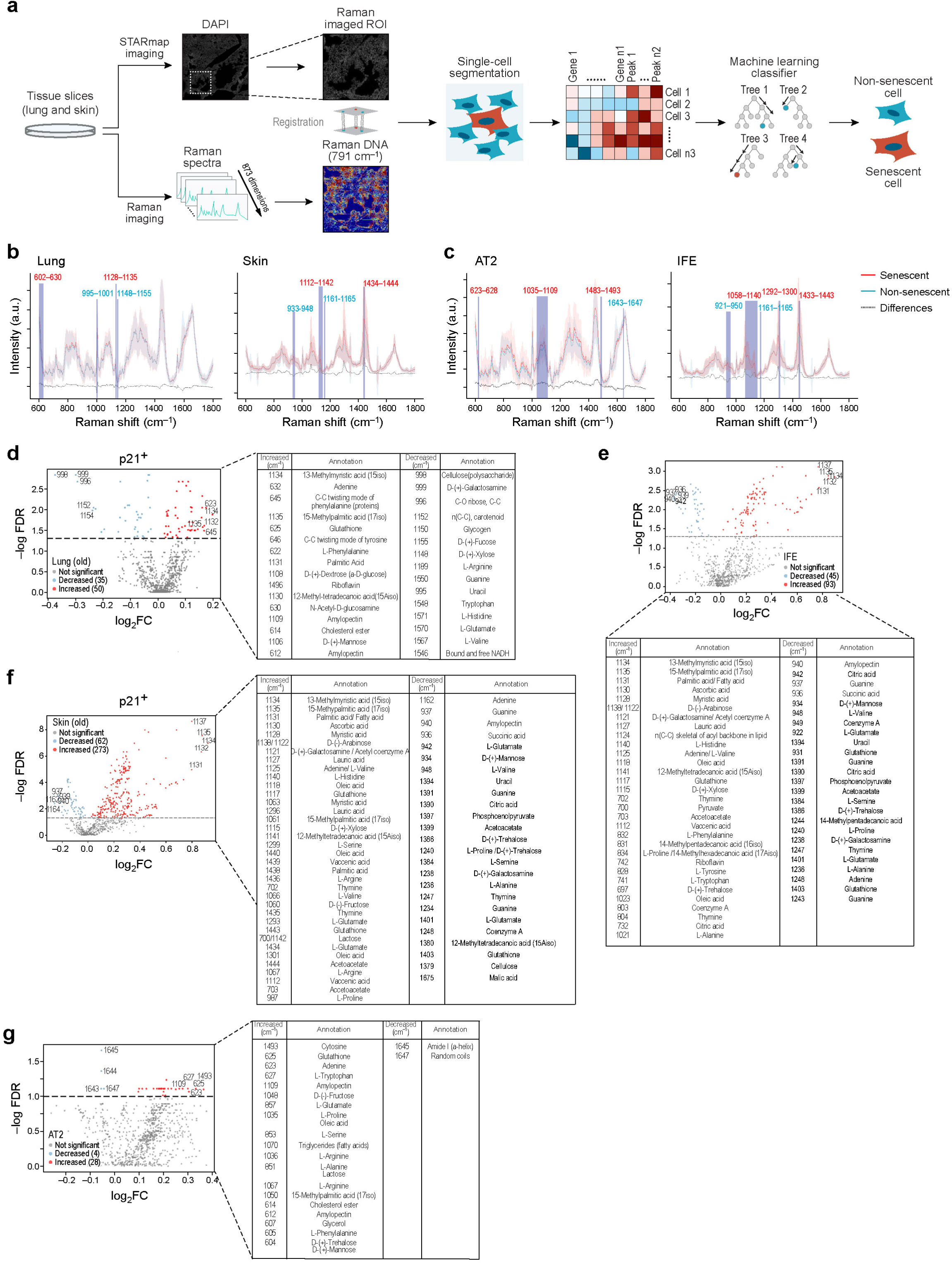
Raman spectral profiling of senescent cells across tissues at single-cell resolution. **a**, Workflow for integrating Raman imaging and STARmap-ISS for senescent cell identification. Raman spectra data (873 dimensions) and STARmap-ISS transcriptomic data (890 genes) were obtained from the same tissue slice. Raman DNA signal (791 cm⁻¹) and DAPI signal (from STARmap-ISS) were used for single-cell registration and segmentation. This integration allowed for the construction of RamanOmics, combining both transcriptomic and Raman measurements within individual segmented cells. Using a custom machine learning classifier, cells were classified into senescent and non-senescent categories. **b-c**, Comparison of the normalized Raman spectra (600-1800 cm⁻¹) of senescent cells (red line) and non-senescent cells (blue line) in mouse lung (**b**, left), mouse skin (**b**, right), AT2 cells (**c**, left), and IFE cells (**c**, right). Dashed lines represent the differential spectrum, indicating variations between senescent and non-senescent cells. Shaded regions with Raman shift values indicate areas with significant DPs. Red numbers denote clusters enriched for increased DPs, blue numbers denote clusters enriched for decreased DPs. **d-g**, Volcano plots showing increased and decreased DPs with corresponding biological annotations in senescent cells from mouse lung (**d**), AT2 cells (**g**), mouse skin (**f**) and IFE cells (**e**) in old samples.

After performing cell type label transfer, we identified 16 cell types in lung and 20 cell types in skin from the aligned cells between STARmap-ISS and Raman imaging (**Extended data Fig. 8a; Table S2**). They represent the majority of cell types detected in our snRNA-seq data with a consistent pattern of senescent cell proportions observed across different samples and age groups from snRNA-seq (**Fig. 2a, b and Extended data Fig. 8b**). These results highlight the robustness and accuracy of our integrated multimodal approach in characterizing the cellular landscape across tissues.

To localize senescent cells within Raman images, we used *p21^+^* cells identified from STARmap-ISS as spatial anchors. This enabled us to pinpoint the corresponding locations and boundaries of these senescent cells in the Raman images (**Fig. 4a**). The subcellular resolution of Raman imaging allowed us to extract precise Raman spectra for both senescent and non-senescent cells via Raman-STARmap image registration (**Fig. 4b, c and Extended data Fig. 9a, b; Methods**), providing detailed insights into the biochemical compositions of these cells.

Next, we compared Raman spectral intensities between old and young cells, as well as between senescent and non-senescent cells across different cell types. We focused on the fingerprint region of the Raman spectra (600-1800 cm⁻¹, 873 dimensions in total), corresponding to key biomolecules such as proteins (amide I, 1640-1680 cm⁻¹; amide III, 1230-1310 cm⁻¹), lipids (1400-1500 cm⁻¹, 1250-1300 cm⁻¹, 1200-1050 cm⁻¹), and nucleic acids (guanine, 785 cm⁻¹, 937 cm⁻¹ and 1234 cm⁻¹; adenine, 536 cm⁻¹, 1125 cm⁻¹ and 1482 cm⁻¹; cytosine, 792 cm⁻¹ and 1275 cm⁻¹)^78,79^. At single-peak resolution, we visualized the full spectral distributions at both the tissue and cell type levels for old versus young cells, and they shared at multiple differential peaks (DP) regions, which reflect unique chemical bonds and vibrational modes of biomolecules in cells (**Extended data Fig. 9a, b**). In parallel, we examined *p21*^+^ and *p21*^-^ cells in mouse lung and skin, both at tissue and cell type resolved manner, and identified several highly variable peak regions indicative of biochemical alterations associated with senescence (**Fig. 4b, c**).

To quantify these differences at the single-cell level, we averaged Raman spectral intensity values from pixels within segmented single cells, aggregating spectral intensities over defined cell boundaries to obtain single-cell Raman spectra (**Methods**). This allowed us to compare Raman intensities between different cells at single-cell resolution.

At the tissue level, old lung samples exhibited higher Raman intensity than young samples (46.7208±28.5472 vs. 43.2734±11.3974, p= 2.10E-13), whereas Raman intensity was lower in old T cells (42.4769±20.1176 vs. 43.0874±11.1290, p= 4.60E-07), although this difference was not significant in skin at either the tissue or cell type level (**Extended data Fig. 10a, c**). Interestingly, global *p21^+^* senescent cells (42.4222±21.1448 vs. 45.4257±11.0594, p= 1.36E-03) and *p21^+^*senescent MES (33.9772±6.5056 vs. 50.7784±11.3752, p= 1.59E-02) showed decreased Raman intensity in old compared to young lung samples (**Extended data Fig. 10b; Table S3**). In contrast, mouse skin exhibited increased Raman intensity in global *p21^+^*senescent cells and *p21^+^*IFE (103.8386±92.1990 vs. 75.2866±72.7948, p= 3.15E-03) in young samples (**Extended data Fig. 10d; Table S3**). Together, these results indicate that aging in lung is generally associated with elevated Raman intensity at the tissue level but reduced intensity in specific immune population, whereas in skin, cell type specific spectral profiles highlight distinct metabolic adaptations and functional states that differentiate aging from senescence.

Leveraging the broad wavelength range (600-1800 cm^-1^) captured from hyperspectral Raman imaging, we investigated biochemical differences across tissues and cell types in aging and senescence analysis. A total of 360 decreased and 422 increased DPs were identified between young and old mouse lung tissues (with adjusted p≤ 0.05), while 462 decreased and 276 increased DPs were found in old versus young skin (**Extended data Fig. 9c, d**). Within Endo of the lung, 247 decreased and 267 increased DPs were detected, largely mirroring the tissue-level lung changes (**Extended data Fig. 9e**). Similarly, in the skin, 441 decreased and 277 increased DPs were observed in IFE cells, highly consistent with the tissue-level Raman shifts (**Extended data Fig. 9f**).

In the senescence analysis, we identified 50 increased and 35 decreased DPs in *p21*⁺ senescent cells compared to *p21*⁻ non-senescent cells in old lung (**Fig. 4d; Table S4**), and 273 increased and 62 decreased DPs at the tissue level in old skin (**Fig. 4f; Table S4**). Cell type specific analysis further revealed 28 increased and 4 decreased DPs in senescent AT2 cells of old lung (**Fig. 4g**), and 93 increased and 45 decreased DPs in senescent IFE cells of old skin (**Fig. 4e**). These results show strong concordance in Raman DPs between tissue-level and cell-type specific analyses. In the old vs. young comparisons, endothelial cells mirror the whole-lung profile, and IFE mirrors whole-skin (**Extended data Fig. 9c-f**). Likewise, in the senescent vs. non-senescent analyses, senescent AT2 cells track with senescent cells in old lung, and senescent IFE cells track with senescent cells in old skin (**Fig. 4d, e**). Together, this overlap underscores the robustness of these peak shifts in capturing aging-and senescence-associated biochemical remodeling at both bulk and single-cell resolution.

Our analysis revealed consistent Raman spectral changes associated with aging and senescence in mouse lung and skin tissues. Leveraging on the Raman peak annotation database^80,81^, we observed in old lung tissue and Endo a significant increase in the 1145/996 cm⁻¹ peaks (linked to “citric acid” and “C-O ribose, C-C”), while 782/790 cm⁻¹ peaks (associated with nucleic acids) were reduced (**Extended data Fig. 9c, e**). In the old skin tissue and IFE cell type, we observed upregulation of 1440/1438 cm⁻¹ peaks, corresponding to trilinolein and myristic acid, alongside reductions in 1524/1517 cm⁻¹ peaks linked to carotenoids and L-glutamate, highlighting the increased lipid and fatty acid signatures in old tissues (**Extended data Fig. 9d, f**). Peaks at 1134/1135 cm⁻¹, attributed to branched-chained fatty acids (BCFAs) such as 13-methylpalmitic acid (15iso) and 15-methylpalmitic acid (17iso), were significantly elevated in *p21^+^* senescent cells across tissue types and specifically IFE cells (**Fig. 4d-f**). BCFAs’ unique structural and functional properties are known to influence membrane fluidity, cellular signaling, inflammation, and metabolic responses^82^. In senescent AT2 cells, increased peaks at 1493/623 cm⁻¹ were observed, corresponding to cytosine and adenine (**Fig. 4g**), suggesting alterations in nucleotide metabolism. Conversely, saccharide-associated peaks at 996/999 cm⁻¹, linked to compounds like ribose (monosaccharide) and D-(+)-galactosamine, a major component of glycoproteins, and peaks at 1162/769 cm⁻¹, related to adenine and amylopectin, showed marked reductions in old mouse lung and skin, respectively (**Fig. 4d, f**). At the cell type level, spectral declines were also noted, with peaks at 1645/1647 cm⁻¹ (amide I and random coils) decreased in senescent AT2 cells (**Fig. 4g**), and peaks at 940/937 cm⁻¹ (Amylopectin and Guanine) reduced in senescent IFE cells (**Fig. 4e**). These changes indicate metabolic reprogramming in senescent cells, marked by increased lipid accumulation, reduced carbohydrate metabolism, and altered protein and nucleic acid structures, collectively contributing to the senescent phenotype.

To spatially visualize Raman intensities within STARmap-ISS segmented cells, we mapped Raman signals onto these segmented cells (**Methods**). Consistent with our previous DP analysis, we observed notable changes in Raman intensities (both increases and decreases) between senescent and surrounding non-senescent cells (**Extended data Fig. 11a, b**), especially within the most variable peak regions at the subcellular level. Specifically, the peaks associated with fatty acid at 1134 cm^-1^ (13-methylpalmitic acid (15iso)), 1135 cm⁻¹ (15-methylpalmitic acid (17iso)) exhibited elevated intensities in senescent cells across lung and skin tissue, as well as in senescent IFE cells at the cell type level (**Extended data Fig. 11a, b**). However, senescent AT2 cells displayed distinct peak changes, with nucleic-acid-associated peaks at 1493 cm^-1^ (cytosine) and 625 cm⁻¹ (adenine) showing increased Raman intensities (**Extended data Fig. 11b**). Conversely, Raman intensities decreased at various peaks at both global tissue and cell type levels. Peaks associated with saccharides at 999 cm⁻¹ (D-(+)-galactosamine) and 934 cm^-1^ (D-(+)-mannose) showed reduced intensities in senescent cells in lung and skin tissues, respectively (**Extended data Fig. 11a**), while intensities at 1643 cm⁻¹ and 1645 cm⁻¹ (Amide I (a-helix)) in senescent AT2 cells and peaks at 940 cm⁻¹ (amylopectin) and 937 cm⁻¹ (guanine) in senescent IFE cells were reduced (**Extended data Fig. 11b**)^80,81^. These Raman intensity changes at tissue-wide and cell type-specific levels underscore the biochemical alterations of cellular senescence and reveal potential Raman markers to understand cellular senescence.

Given that the DPs contribute significantly to distinguishing old cells to young cells, as well as senescent cells from non-senescent cells, we developed a DP barcode system to assign each cell a unique barcode ID for accurate classification of senescence based on Raman features (**Methods**). By integrating the top 30 increased and top 30 decreased DPs (top 4 increased and top 28 decreased DPs for AT2 cells), we generated specific barcodes for old cells and young cells (**Extended data Fig. 12a, b**), as well as for *p21^+^* senescent and *p21^-^* non-senescent cells across different tissues and cell types, as well as for *p21^+^* senescent cells from both old and young samples (**Extended data Fig. 12c, d**). This barcode visualization clearly delineates the DP differences between old and young, senescent and non-senescent cells, providing a robust and unique framework for tracking and comparing senescence-associated biomolecule alterations across different conditions. This approach enables us to identify unique and overlapping biochemical signatures that define senescent cell states, offering new insights into the heterogeneity and functional consequences of cellular senescence in aging tissues.

### Spatially resolved multimodal molecular and biochemical landscapes of senescence

Anchoring on *p21*⁺ cells allow us to directly connect transcriptomic and biochemical alterations to the senescence phenotype. Using this framework, we applied a multimodal approach integrating Raman spectra with snRNA-seq data in old samples, with STARmap expression profiles serving as spatial anchors (**Methods**). From the aligned cells, we selected the top DPs (top 30 increased and top 30 decreased for tissue level and IFE cells, 28 increased and 4 decreased for AT2 cells) and combined with all selected differentially expressed genes (DEGs) from snRNA-seq (37 upregulated and 29 downregulated for mouse lung, 17 upregulated and 29 downregulated for mouse skin, 7 upregulated and 2 downregulated for AT2 cells, 5 upregulated and 3 downregulated for IFE cells) (**Table S6**) to create a multimodal digital representation of cells with three distinct modalities (**Fig. 1a**). To evaluate the performance of the combined multimodal features in identifying senescent cells, we built a random forest classifier based on individual Raman features, snRNA-seq features, and combined multimodal features using 70% of the cells for training and 30% for testing to compare the performance in classifying senescence cells. Our results demonstrated that the performance metrics (Accuracy, AUC, and Precision) significantly improved in both mouse lung and skin samples when Raman features were added to the snRNA-seq data, with Accuracy increased by 5.36% to 5.87% (from 0.7368 to 0.7763 for mouse lung, and from 0.6182 to 0.6545 for mouse skin), and AUC score increased by 2.64% to 5.91% (from 0.7450 to 0.7647 for lung and from 0.6819 to 0.7222 for skin) and Precision score increased by 5.36% to 13.45% (from 0.6809 to 0.7174 for lung and from 0.6296 to 0.7143 for skin) respectively (**Fig. 5a**). We also observed Raman-only model in skin exhibited superior predictive performance relative to Raman-only model in lung across all evaluated metrics (e.g., 0.6000 vs. 0.4868 for Accuracy, 0.672 vs. 0.5367 for AUC). This enhanced predictive power for skin may correspond with findings from snRNA and STARmap analyses to show a higher proportion of *p21^+^* senescent cells in skin tissue (**Fig. 3e, f**). As the body’s outermost barrier, skin is continuously exposed to environmental stressors, such as UV radiation and pollution, which likely accelerate senescence and induce pronounced biochemical changes detectable by Raman spectroscopy.

**Figure 5.**
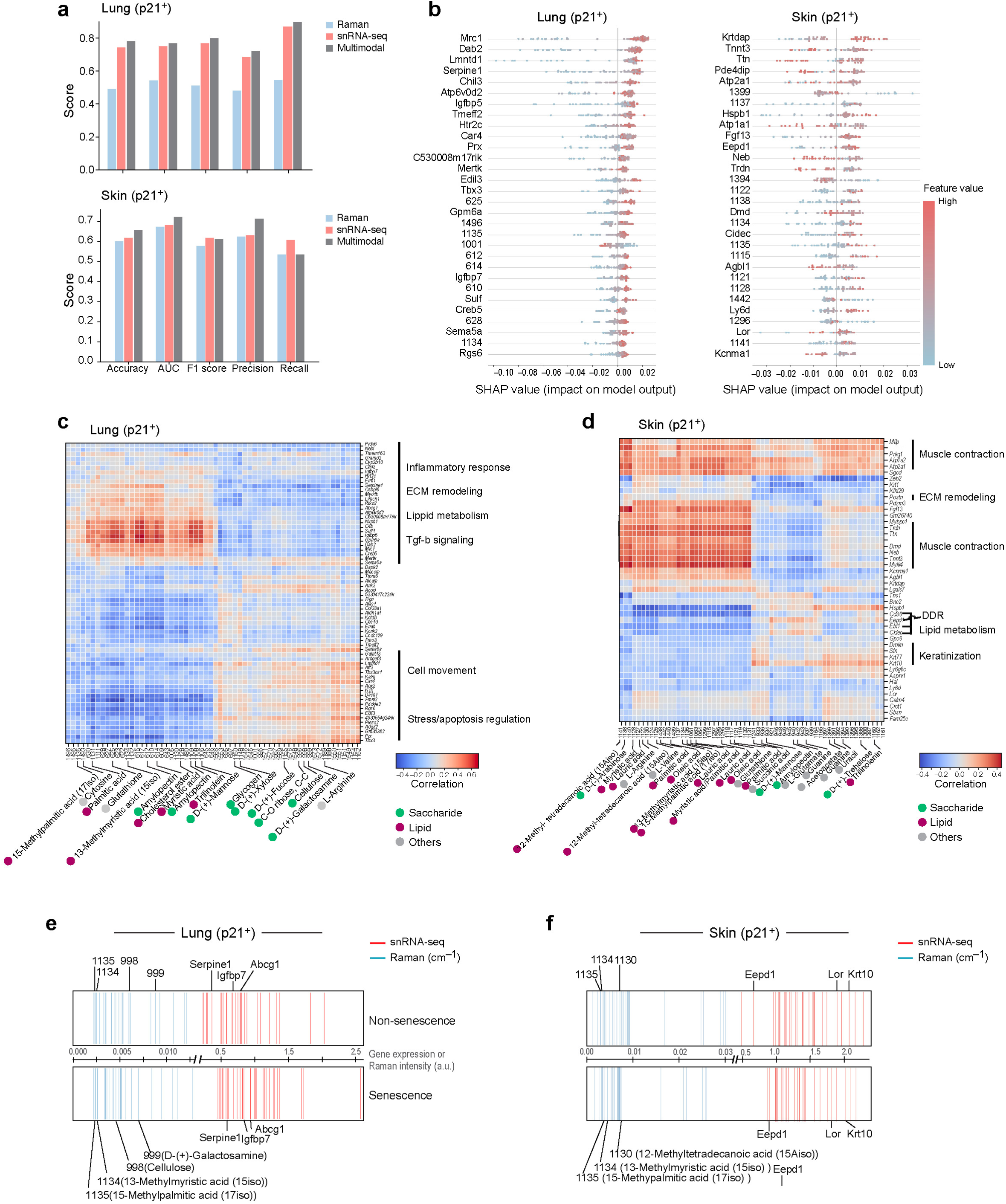
Multimodal integration and characterization of senescent cells. **a**, Bar graph displaying the classification performance of different metrics using Raman, snRNA-seq or integrated multimodal datasets from mouse lung (top) or mouse skin (bottom). **b**, SHAP summary dot plots showing the impact of top 30 important features (Raman peaks and genes) on senescent cell prediction at single-cell level. X-axis represents the SHAP value that indicates senescent cell predicting power of each feature, Y-axis lists the features sorted by importance. Each dot indicates a single cell. **c-d**, Heatmap showing the correlations between the selected top DPs (60 for lung, 60 for skin) and DEGs (66 for lung, 46 for skin) in *p21*^+^ senescent cells from mouse lung (**c**) and mouse skin (**d**). Raman peaks and gene functions were manually annotated. **e-f**, Stripe barcodes representing the most important multimodal features identified by the random forest in *p21*^+^ senescent cells versus *p21*^-^ non-senescent cells in mouse lung (**e**) and mouse skin (**f**). The barcodes display the top 60 identified features (32 DEGs and 28 DPs for mouse lung and 30 DEGs and 30 DPs for mouse skin). Raman peaks arranged based on average peak intensities with representative DPs annotated for their biological meanings. Genes arranged based on average gene expression levels with representative genes highlighted.

We then extracted and ranked the top 30 multimodal features (DPs and DEGs) that significantly contributed to the prediction using both random forest feature importance and SHapley Additive exPlanations (SHAP) methods (**Fig. 5b and Extended data Fig. 13a, b**). These two approaches yielded largely consistent results, as they identified many of the same genes and Raman peaks as key contributors to prediction accuracy. Specifically, in agreement with our previous snRNA-seq and Raman imaging findings, we identified the DEGs *Mrc1*, *Dab2*, *Igfbp7*, and *Serpine1,* and DPs 1134 cm^-1^ (13-methylpalmitic acid (15iso)), 1135 cm⁻¹ (15-methylpalmitic acid (17iso)), and 625 cm⁻¹ (adenine) from mouse lung, as well as genes *Hspb1*, *Lor*, and *Krtdap,* and peaks 1134 cm^-1^ (13-methylpalmitic acid (15iso)), 1135 cm⁻¹ (15-methylpalmitic acid (17iso)), and 1137 cm⁻¹ from mouse skin, which were upregulated in senescent cells and demonstrated strong predictive power in both methods (**Fig. 5b and Extended data Fig. 13a**). These findings suggest that by integrating multimodal data, including molecular and vibrational–biochemical features, it significantly improves the accuracy of senescent cell identification and the robustness of our feature importance extraction methods.

Subsequently, we analyzed the positive and negative contributions of the top-ranked features in identifying senescent cells using both feature importances from the classifier and SHAP values (**Methods**). In mouse lung, Raman peak at 625 cm⁻¹ (adenine) and genes *Mrc1*, *Dab2,* and *Serpine1* consistently ranked among the most important features in both senescent and non-senescent cells (**Extended data Fig. 13c**).

Additionally, specific Raman peaks, such as 612 cm⁻¹ (amylopectin), were distinctly upregulated in senescent cells in mouse lung tissues, contributing significantly to their classification (**Extended data Fig. 13c**). In mouse skin, *Krtdap* and *Dmd* were among the top-ranked markers of senescent cells, while *Ttn* and *Cidec* were preferentially ranked in non-senescent populations (**Extended data Fig. 13d**). Key lipid-associated Raman peaks, 1134 cm⁻¹ (13-methylpalmitic acid (15iso)), and 1135 cm⁻¹ (15-methylpalmitic acid (17iso)), were among the most important discriminative features in both classes (**Extended data Fig. 13d**). Taken together, these results indicate that transcriptomic features, such as *Dab2* and *Serpine1* in lung while *Ttn* and *Krtdap* in skin, are critical for distinguishing senescent from non-senescent cells, whereas Raman-derived biochemical features provide complementary and orthogonal information that enhances the precision and robustness of senescence classification.

To further explore which features have a positive or negative influence on prediction, we visualized SHAP values at the single-cell level, highlighting key features influencing the model’s predictions. In mouse lung, genes *Dab2*, *Igfbp7*, and *Serpine1*, and Raman peaks at 625 cm⁻¹ (adenine) and 1135 cm⁻¹ (15-methylpalmitic acid (17iso)) positively contributed to senescent cell identification (red dots with positive SHAP values), while peaks at 1001 cm⁻¹ was associated with non-senescent cells (red dots with negative SHAP values) (**Fig. 5b**). Similarly, in mouse skin, genes *Krtdap*, *Hspb1*, and *Lor,* and peaks at 1134 cm⁻¹ (13-methylpalmitic acid (15iso)), 1135 cm⁻¹ (15-methylpalmitic acid (17iso)) and 1128 cm⁻¹ (myristic acid) supported senescent cell identification (**Fig. 5b**). These findings align closely with our independent snRNA-seq and Raman analyses, demonstrating the consistency and robustness of our multimodal approach for accurately identifying senescent cells across different tissues.

### Intertwined and multilayered transcriptional, metabolic, and structural rewiring in senescence

By directly linking Raman spectral peaks to individual genes, we can uncover multilayered molecular pathways and biochemical compositions that define the senescent phenotype. To achieve this, we performed correlation analyses between DPs and DEGs which enable us to assign clear biological significance to specific Raman features (**Fig. 1a; Methods**).

In mouse lung, we identified distinct Raman peak clusters, including 602-630 cm⁻¹ (607 cm⁻¹, glycerol; 614 cm⁻¹, cholesterol ester), 1106-1111 cm⁻¹ (1108 cm⁻¹, α-D-glucose; 1109 cm⁻¹, amylopectin), and 1128-1135 cm⁻¹ (1130 cm⁻¹, 12-methyl-tetradecanoic acid (15Aiso); 1131 cm⁻¹, palmitic acid; 1134 cm⁻¹, 13-methylmyristic acid (15iso); 1135 cm⁻¹, and 15-methylpalmitic acid (17iso)). These peaks were significantly elevated in senescent cells and correlated with upregulation of genes involved in ECM remodeling (*Serpine1*, *Sulf1*), lipid metabolism (*Abcg1*), and TGF-b signaling pathway (*Dab2*, *Igfbp7*) (**Fig. 5c**). In contrast, peak clusters 995-1001 cm⁻¹ (996 cm⁻¹, C-O ribose, C-C; 999 cm⁻¹, D-(+)-galactosamine), 1148-1155 cm⁻¹ (1148 cm⁻¹, D-(+)-xylose; 1150 cm⁻¹, glycogen; 1152 cm⁻¹, n(C-C), carotenoid; 1155 cm⁻¹, D-(+)-fucose), and 1544-1576 cm⁻¹ (1550 cm⁻¹, guanine; 1567 cm⁻¹, L-valine; 1570 cm⁻¹, L-glutamate), which were reduced in senescent cells, were associated with anti-apoptotic activity (*Tbx3*)^83^ and cell motility (*Fmnl2*, *Kalrn*)^84,85^ (**Fig. 5c**). Collectively, these findings highlight key biochemical alterations in senescent cells, particularly in ECM remodeling, lipid metabolism, and stress response pathways, which together contribute to the senescence phenotype in lung tissue.

In mouse skin, elevated Raman peak clusters at 1112-1141 cm⁻¹ (e.g., 1128 cm⁻¹, myristic acid; 1130 cm⁻¹, 12-methyltetradecanoic acid (15Aiso); 1131 cm⁻¹, palmitic acid and other fatty acids; 1134 cm⁻¹, 13-methylmyristic acid (15iso); 1135 cm⁻¹, 15-methylpalmitic acid (17iso)) and at 1434-1444 cm⁻¹ (e.g., 1438 cm⁻¹, palmitic acid; 1439 cm⁻¹, vaccenic acid; 1440 cm⁻¹, oleic acid), coinciding with transcriptomic downregulation of genes involved in ECM (*Postn*) and muscle contraction (*Tnnt3*, *Ttn*)^86,87^ and upregulation of keratinization genes (*Sbsn*, *Lor*)^88^ (**Fig. 5d**). In contrast, decreased peaks at 933-948 cm⁻¹ (e.g., 934 cm⁻¹, D-(+)-mannose; 940 cm⁻¹, amylopectin) and 1161-1165 cm⁻¹ (e.g., 1161 cm⁻¹, quinoid ring deformation; 1162 cm⁻¹, adenine) were associated with downregulation of DNA damage response gene *Eepd1*, and upregulation of skin barrier-maintenance genes (*Sfn*, *Krt10*). Together, these multimodal signatures indicate that senescent cells undergo pronounced lipid remodeling and reinforcement of differentiation programs, accompanied by impaired DNA repair and compensatory barrier homeostasis, hallmarks of skin aging and senescence.

Subsequently, we further analyzed the DP-DEG correlation at the cell type level for AT2 and IFE. In senescent AT2 cells in lung, elevated Raman peak clusters at 600-615 cm⁻¹ (e.g., 607 cm⁻¹, glycerol; 614 cm⁻¹, cholesterol ester), and 623-628 cm⁻¹ (e.g., 623 cm⁻¹, adenine; 625 cm⁻¹, glutathione), were associated with genes *Pcsk6 and Serpine1* (ECM remodeling) and *Meg3* (Cell senescence). By contrast, decreased peaks in the 1643-1647 cm⁻¹ range were linked to the downregulation of *Tex11* (DNA damage repair) and *Acoxl* (lipid metabolism), indicating impaired DNA repair capacity and reduced lipid metabolic activity (**Extended data Fig. 13e**). These changes suggest a progressive decline in genome maintenance and metabolic flexibility, reflecting a shift toward terminally arrested, fully senescent AT2 cells.

In skin senescent IFE cells, increased peaks at 1112-1141 cm⁻¹ (e.g., 1131 cm⁻¹, palmitic acid; 1135 cm⁻¹, 15-methylpalmitic acid (17iso)) correlated with downregulated genes *Rad51b* (DNA damage repair)^89^ and *Sema5a* (inflammatory response)^90^, suggesting that biochemical lipid remodeling in senescent cells occurs in the context of impaired DNA repair capacity and attenuated immune regulatory signaling. In contrast, decreased Raman peaks at 922-949 cm⁻¹ (e.g., 934 cm⁻¹, D-(+)-mannose) and 1382-1404 cm⁻¹ (e.g., 1392/1395 cm⁻¹, uracil) were associated with the downregulation of *Antxr1* (ECM remodeling)^91^, but strongly correlated with the upregulation of keratinization-related genes (*Krt10*, *Krt77*, *Krtdap*, *Sbsn*, *Dmkn*) (**Extended data Fig. 13f**).

To increase resolution beyond transcript-only markers, we integrated high-dimensional Raman spectra with RNA-seq and extended our DP barcoding to a RamanOmics barcode that incorporates DEGs into DP framework. Using the top predictive multimodal features (**Table S4; Methods**), we generated robust cell type specific barcodes that distinguish *p21*⁺ senescent from *p21*⁻ non-senescent cells across both global tissues (lung, skin) and specific cell types (AT2, IFE) (**Fig. 5e, f and Extended data Fig. 13g, h**).

This approach provides an intuitive, quantitative display of senescence multimodal signatures, enabling precise identification of senescent states. The RamanOmics barcode sharpens senescence classification and resolves cellular heterogeneity across tissues, establishing a broadly applicable bio-metrological framework for measuring and characterizing senescent cells.

### Validation of multimodal senescence programs in wound repair

To validate the newly identified senescent signatures and programs in a pathologically relevant context, we employed a mouse skin wound-healing model, which faithfully recapitulates the transient and dynamic induction of senescent cells during tissue regeneration^92^. Using STARmap-ISH following Raman imaging, we observed that on day 3 post-wounding (D3), expression of the canonical senescence marker *p21* was markedly increased compared with unwounded skin (D0) in old mouse (**Fig. 6a and Extended data Fig. 14**), consistent with prior reports^92^. Importantly, *p21* induction was accompanied by upregulation of potential novel senescence markers (*Krt10*, *Dmkn*, *Lor, Sbsn, Sfn*) (**Fig. 6b and Extended data Fig. 14**), which were identified from our snRNA-seq analysis (**Fig. 2d**). These findings functionally validate our transcriptomic discoveries and establish these genes as bona fide senescence markers during tissue repair.

**Figure 6.**
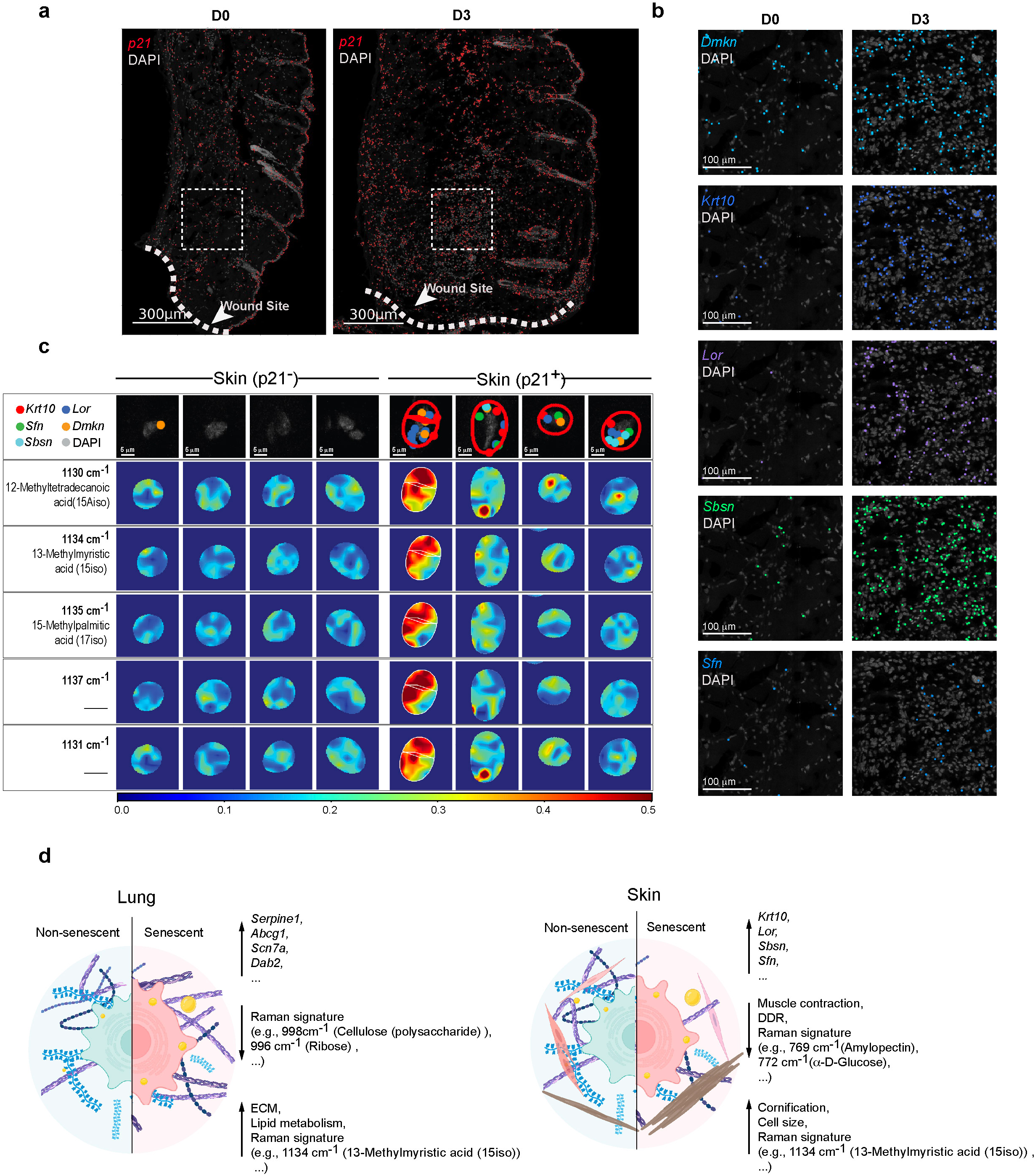
Validation of novel molecular and biochemical signatures in a mouse skin wound-healing model. **a**, Spatial distribution of *p21* expression at the wound site of old mouse skin at baseline (D0) and three days post-injury (D3), as detected by STARmap-ISH. **b**, Higher-magnification views of the regions delineated by the white dashed boxes in (**a**), showing expression of *Dmkn*, *Krt10*, *Lor*, *Sbsn*, and *Sfn* in D0 and D3 sections. **c**, Representative Raman intensity maps of senescence-associated peaks overlaid with subcellular localization of senescence marker genes (*Krt10*, *Sfn*, *Lor*, *Sbsn*, *Dmkn*) detected by STARmap-ISH in *p21*⁻ and *p21*⁺ cells. Red lines indicate the segmented cell boundary of *p21*⁺ cells. **d**, Diagrams illustrating the multimodal features of senescent cells compared to non-senescent cells in mouse lung and skin.

We next examined whether the biochemical markers of senescence identified by Raman imaging were also conserved during wound repair. Consistent with our multimodal analysis, senescent cells in wounded skin displayed increased Raman intensities at 1130/1134/1135 cm⁻¹, corresponding to 12-Methyltetradecanoic acid (15Aiso), 13-Methylmyristic acid (15iso), 15-Methylpalmitic acid (17iso), along with increased expression of the novel senescent markers (**Fig. 6c**). These changes reinforce the conclusion that lipid remodeling is the integral biochemical feature of the senescent state *in vivo*.

Together, these experiments confirmed *Dmkn*, *Krt10*, *Lor*, *Sbsn*, and *Sfn*, along with biochemical markers 12-Methyltetradecanoic acid (15Aiso), 13-Methylmyristic acid (15iso), 15-Methylpalmitic acid (17iso) at Raman peaks 1130/1134/1135cm⁻¹, as novel senescence markers in mouse skin and confirmed their functional relevance during wound repair and regeneration. More broadly, they demonstrate the power of integrating Raman spectroscopy with spatial transcriptomics strengthens validation and mechanistic deconvolution of senescence signatures in complex tissue settings.

In summary, our study demonstrates that senescent cells in mouse lung exhibit increased lipid metabolism (e.g., *Abcg1*), ECM remodeling (e.g., *Serpine1, Sulf1*) and increased lipid-associated Raman peaks (e.g., 1134 cm^-1^ (13-methylmyristic acid (15iso)), 1135 cm^-1^ (15-methylpalmitic acid (17iso)), along with decreased saccharide-associated Raman peak intensities (e.g., 998 cm^-1^ (cellulose(polysaccharide)), 999 cm^-1^ (D-(+)-galactosamine)) (**Fig. 6d**). In contrast, senescent cells in mouse skin show upregulated cornification or terminal differentiation (e.g., *Sfn*, *Krt10*) and lipid-associated Raman peaks (e.g., 1134 cm^-1^ (13-methylmyristic acid (15iso)), 1135 cm^-1^ (15-methylpalmitic acid (17iso))), with reduced muscle contraction (e.g., *Tnnt3*, *Dmd*), DDR (e.g., *Eepd1*) and the saccharide-associated Raman peaks (e.g., 769 cm^-1^ (amylopectin), 772 cm^-1^ (a-D-glucose)) (**Fig. 6d**). This tissue-wide, cell-type-resolved, multimodal framework establishes biochemical-transcriptional couplings of senescence across organs, resolving shared and tissue-specific axes. By bridging biochemical composition with gene expression at single-cell resolution, our approach not only enhances the fidelity of senescence detection but also provides a framework to interrogate how senescent states contribute to diverse biological processes such as wound healing, fibrosis, and tissue remodeling.

## DISCUSSION

Aging reshapes tissues through the accumulation of senescent cells, yet current definitions of senescence still depend largely on transcriptomic or histological markers that fail to capture its underlying biochemical remodeling. To address this gap, we developed RamanOmics, a multimodal framework that integrates single-nucleus transcriptomics, spatial transcriptomics, and label-free Raman imaging (**Fig. 1a**). This approach demonstrates that senescence is characterized not only by molecular rewiring but also by biochemical shifts that are detectable non-destructively *in situ*.

This was evidenced by increased fatty-acid-associated Raman peaks (e.g., 1135/1128 cm⁻¹ in the lung and 1141/1160 cm⁻¹ in the skin) that correlated with the upregulation of the lipid transport gene *Abcg1* in the lung and downregulation of the lipid metabolism gene *Cidec and Cd36* in skin (**Fig. 5c, d**), indicating a conserved, age-related shift in lipid composition across tissues. These lipid alterations were not isolated events but occurred alongside major transcriptional programs: activation of ECM remodeling (*Serpine1*, *Sulf1*) and TGF-β signaling (*Dab2*, *Igfbp7*) in lung, together with downregulation of DNA damage response (*Eepd1*) and activation of keratinization pathways (*Lor*, *Sbsn*) in skin (**Fig. 5c, d**). Together, these findings point to a coupled lipid-matrix-differentiation program that reinforces the pro-fibrotic and metabolically reprogrammed state of senescent cells. At the same time, tissue-specific features became evident: skin showed reductions in nucleic-acid-associated Raman peaks (e.g., 1162 cm⁻¹-adenine, 937 cm⁻¹-guanine), whereas lung displayed decreases in saccharide-and amino-acid-linked Raman peaks (e.g., 998 cm⁻¹-Cellulose, 999 cm⁻¹-D- (+)-galactosamine, 1155 cm⁻¹-D-(+)-fucose, 1189 cm⁻¹-L-arginine). Thus, senescence reflects both a core biochemical program and context-dependent adaptations shaped by the tissue microenvironment.

The integration of molecular and biochemical modalities also revealed unexpected biology. BCFAs enrichment was consistently detected in *p21*⁺ senescent cells across lung and skin, including epidermal keratinocytes (**Fig. 4d, e**). Because BCFAs in mammals may arise from both endogenous remodeling and microbiome or dietary sources, their consistent enrichment across contexts suggests a potential crosstalk between systemic lipid metabolism, tissue niche, and senescence biology. Additionally, Raman mapping revealed subcellular heterogeneity in nucleic acid and lipid features within senescent cells, illustrating how multimodal integration can capture aspects of senescence invisible to transcriptomics alone.

The wound-healing model provided critical pathological validation. Senescent cells are transiently induced during repair and are essential for optimal tissue regeneration, largely through paracrine signaling such as PDGF-AA^93^. By applying RamanOmics to wounded skin, we confirmed transient *p21* induction and validated upregulation of *Sbsn*, *Dmkn*, *Krt10*, *Sfn* and *Lor* accompanied by the previously detected Raman shifts (e.g., increased 1134/1137 cm^-1^ and decreased 1162/937 cm^-1^), relative to *p21*^-^ cells (**Fig. 6a-c**). These results showed that multimodal senescence signatures identified in steady-state aging extend to regenerative contexts, reinforcing their functional relevance.

Like any newly developed technology, RamanOmics faces limitations that also point to opportunities. First, Raman imaging cannot yet capture very large tissue regions at subcellular resolution, restricting our analyses to selected fields of view and sample size. Second, *p21^+^* senescent cells remain a rare population even in old tissues, limiting the number of cells available for training and potentially reducing classifier power. This highlights the need for high-speed Raman microscopy, larger multi-tissue datasets, and integration of consensus senescence signatures. Finally, although spectral assignments for fatty acids and saccharides were supported by references^94^, their precise biological interpretation and annotation require further validation, including isotope tracing and complementary mass spectrometry imaging.

Despite these challenges, the translational implications are significant. Non-destructive Raman features alone could serve as biomarkers of senescence, enabling *in vivo* monitoring of tissue aging or evaluation of senolytic therapies in humans. Coupling biochemical readouts with transcriptomic programs also opens opportunities for high-throughput screening of lipid-pathway modulators and longitudinal tracking of senescence in translational contexts such as skin aging, fibrosis, or wound repair. More broadly, RamanOmics illustrates how multimodal integration can reveal biochemical-genetic couplings that remain hidden to single-modality assays. This modular design also supports integration with additional omics and extension to diverse biological questions, including the characterization of cell states of interest in health and disease.

Together, these data highlight opportunities to extend this framework. Perturbing lipid elongases and branching enzymes, combined with isotope-resolved Raman tracing, will directly test whether lipid remodeling sustains senescence. Expanding to human tissues will establish translational robustness, and applying the framework to fibrosis, regeneration, and cancer will determine whether these biochemical hallmarks are universal or context specific. By uniting spatial transcriptomics with label-free vibrational-biochemical imaging, RamanOmics provides a modular framework for spatiotemporal dissection of aging and a foundation for multimodal profiling of cell states in complex tissues.

## RESOURCE AVAILABILITY

### Lead contact

Further information and requests for resources and reagents should be directed to and will be fulfilled by the lead contact, Dr. Jian Shu

### Materials availability

This study did not generate new unique reagents.

### Data and code availability

All data related to this study are available at: https://data.sennetconsortium.org/ Group: TDA-Massachusetts Institute of Technology

### Code availability

Code for comprehensive analysis as described here, and figure generation as shown here can be found at https://github.com/jian-shu-lab/RamanOmics

## ACKNOWLEGEMENTS

We thank lab members in the Shu lab, the So/Kang lab, the Dou lab, the Phatnani lab for insightful discussions. We thank Leslie Gaffney for editing of the figures. This research is supported by the NIH Common Fund, through the Office of Strategic Coordination/Office of the NIH Director under awards UG3CA275687, UH3CA275687 (P.T.S., J.W.K., J.S.), U54AG076040 (H.P), UG3CA268117, UH3CA268117 (Z.X.D.), NIH New Innovator Award 1DP2AG080320-01 (J.S.), P41EB015871 (J.W.K., P.T.S.), R01DC021326 (J.W.K.), National Cancer Center, Korea (NCC-24H1170) (J.W.K., P.T.S.), Massachusetts General Hospital (J.S.).

## AUTHOR CONTRIBUTIONS

J.S. conceived, designed and directed the study. J.S., P.T.C.S. and J.W.K. co-supervised the collaborative project. Z.D. K.H. and Y.X. bred and fed the young/old mice, generated the wound-healing model and provided valuable discussions. K.Z. and C.C., generated single-nucleus sequencing data with contributions from S.B. and F.V. J.W.K and S.S performed Raman imaging with contributions from M.T., K.Z and K.J.K.K. K.Z. generated STARmap-ISS and STARmap-ISH data with contributions from S.B., S.S., J.Z. and J.Q. F.M. and H.H analyzed snRNA-seq data with contribution from Y.Q., W.Y., and S.D. F.M and H.H analyzed STARmap-ISS and STARmap-ISH data with input from X.C and K.Z. X.C analyzed Raman imaging data and performed integration analysis of snRNA-seq data and Raman imaging data with contribution from F.M, H.H and K.Z. H.P., Z.D. and P.T.C.S provided conceptual, methodological suggestions and feedback. K.Z. assembled figures with contribution from S.S., J.Z., and with input from all authors. K.Z. and J.S. wrote the manuscript with contributions from all authors. All authors read and accepted the manuscript.

## Methods

### Ethical statement

All experiments performed in this study were in accordance with the relevant animal husbandry standards of the Committee on Animal Care.

### Experimental mouse model

p16-3MR mice were generated from Jackson lab (RRID: IMSR_JAX:037045), and skin and lung samples were collected from mice at 2 months (n= 3) and 26 months (n= 3) old.

### Mouse wound-healing model

Full-thickness excisional wounds were generated on the dorsal skin of 2-month-old and 24-month-old p16-3MR mice. Prior to surgery, mice were anesthetized with 3% isoflurane via nose cone and administered subcutaneous analgesia using Ethiqa XR (3.25 mg/kg, 1.3 mg/mL) 30 minutes before wounding. The dorsal area was first shaved with electric clippers and then depilated using a chemical depilatory cream, which was applied for no more than 30 seconds and thoroughly removed with moist gauze to avoid skin irritation. A single 6-mm full-thickness wound, including panniculus carnosus, was created on the central dorsum using a sterile biopsy punch. For each mouse, two samples were collected: (1) the excised wound tissue at day 0, and (2) wound-edge skin at day 3 post-injury using a 6-mm punch biopsy centered at the healing margin. All procedures were conducted in compliance with institutional animal care and use guidelines.

### Tissue cryopreservation

Dissected mouse skin and lung samples were placed into OCT-containing plastic molds, frozen on dry ice, and stored at -80°C.

### Single-nuclei isolation

Single nuclei isolation was conducted according to the 10x Genomics guide (CG000505) using Chromium Nuclei Isolation Kit with RNase Inhibitor (PN-1000494). Briefly, OCT embedded mouse lung and skin tissues were cut into 100 μm slices, and nuclei were isolated from six slices. Lysis buffer was added to the tissue slices, and the tissues were dissociated with a plastic pestle until homogeneous. The mixture was then incubated on ice for 10 minutes. The dissociated nuclei suspension was passed through a nuclei isolation column by centrifugation. Nuclei were pelleted by centrifugation at 500 rcf at 4°C for 3 minutes, followed by the removal of cell debris. The nuclei pellet was washed and resuspended in wash and resuspension buffer, and the suspended nuclei were kept on ice before loading into the 10x Chromium Controller.

### Single-nuclei RNA expression library preparation and sequencing

Isolated nuclei were processed, and single-nuclei libraries were prepared using the Chromium Next GEM Single Cell 3’ Reagent Kits v3.1 (Dual Index) protocol (10X Genomics), loading 8,000 nuclei per sample. Pooled libraries were then sequenced using NextSeq High Output Cartridge kits and a NextSeq 550 sequencer (Illumina).

RNA Libraries were sequenced using the configuration of R1: 28 cycles, R2: 44 cycles, Index1: 10 cycles, Index2: 10 cycles.

### Single-cell RNA-seq data processing

The count matrices obtained from Cell Ranger^95^ were uploaded into R (v4.2.3) to remove the ambient RNA contamination using the SoupX package (v1.6.2)^96^, transforming the resulted count matrices into integer (roundToInt= TRUE), and other parameters as default value.

The resulting matrices were used as input to construct a Seurat object using the Seurat package (v4.3.0)^97^ (using min.cells= 5, and other parameters as default value).

The identification of putative multiplets was done using the scDblFinder (v1.12.0)^94^ (with default parameters) and the DoubletFinder (v2.0.4)^98^ packages (using number of expected doublets (nExp) 5% for all the datasets, and other parameters as default value). Cells labeled as multiplets by any of the two methods were removed from the count matrices.

The Seurat package was used for further filtering low-quality cells. The filtering method was used to detect outliers and cells expressing a high number of mitochondrial transcripts. Only cells with nFeature_RNA, nCount_RNA>300, nFeature_RNA<+2 Standard Deviation from mean, and with percentage of reads mapped to the mitochondrial genome <10% were used for the downstream analysis. A total of 35474 high quality cells was obtained for lung samples and a total of 12128 high quality cells was obtained for skin samples.

The resulted Seurat objects were merged, normalized (using logNormalize as normalization method), used for the identification of the highly variable genes (nfeatures= 3000). The ScaleData and RunPCA functions were applied.

Principal components explaining more than 90% of the data variance were used for uniform manifold approximation and projection analysis. Clustering parameter resolution was set to 0.8 for the function FindClusters of the Seurat package. For the identification of DEGs between clusters, the FindAllMarkers function of the Seurat package was used (with min.pct= 0.25, only.pos= TRUE, and other parameters as default value).

### Cell annotation

For annotating the cell types, a combination of manual and automatic approaches was employed. For mouse lung data, label transfer was performed using Seurat and the mouse lung atlas by Angelidis *et al*.^99^ as the reference. Subsequently, manual annotation was used for the validation and refinement of label transfer results. The DEGs between clusters obtained from Seurat FindAllMarkers function were compared against known cell-type–associated genes. For mouse skin data, label transfer was performed using Seurat and the mouse skin atlas Joost et al.^100^ as the reference. Red blood cells were removed from the reference dataset. Subsequently, manual annotation was used for the validation and refinement of label transfer results. The DEGs between clusters obtained from Seurat FindAllMarkers function were evaluated using established marker-gene profiles.

### Differential gene expression analysis and gene set enrichment analyses

For the identification of DEGs between senescent and non-senescent cells within each condition, and between old and young senescent cells, the FindMarkers function of the Seurat package was used (with logfc.threshold= 0.1, min.pct= 0.01, test.use= “wilcox”, and other parameters as default value). Only genes with |log2FC|>1 and Bonferroni adjusted p<0.05 were considered significant DEGs.

For the identification of DEGs between old and young cells, the FindMarkers function of the Seurat package was used (with logfc.threshold= 0.25, min.pct= 0.25, test.use= “wilcox”, and other parameters as default value). Only genes with |log2FC|>1 and Bonferroni adjusted p<0.05 were considered significant DEGs.

The Gene Ontology (GO) enrichment analysis was performed on significant DEGs using clusterProfiler (v4.6.2)^101^. Enriched GO terms were calculated using enrichGO function (with ont= “BP”, OrgDb= “org.Mm.eg.db”, qvalueCutoff= 0.05, and other parameters as default value).

The pathway enrichment analysis was performed on significant DEGs using gprofiler2 (v0.2.3)^102^. Functional enrichment of gene lists was calculated using gost function (with organism= “mmusculus”, correction_method= “fdr”, domain_scope= “annotated”, sources= c(“KEGG”, “REAC”, “WP”)).

### Cell-cell communication analysis

For inferring the cell-cell communications among cell types from senescent and non-senescent cells, CellChat^103^ (v2.1.2) was used. Cell groups are defined using a combination of senescence status and cell type. Communication probability was computed using population.size= TRUE. Only cell-cell communications found in more than 10 cells in each group were considered for both senescent and non-senescent cells.

### Cell-type composition analysis

Cell-type composition analysis was performed using the R package scProportionTest^104^ Only cell types with FDR<0.01 and log2FC>1 between old and young samples were considered significant.

### STARmap and Raman imaging sample preparation

A quartz-bottom culture dish was coated with a poly-D-lysine solution. OCT-embedded mouse lung and skin tissues were sectioned into 14 μm slices, then fixed with 4% PFA in PBS for 15 minutes, followed by three washes with PBSR (PBS + RNase Inhibitor). Fixed samples submerged with PBSR underwent Raman imaging before being permeabilized with pre-chilled methanol at -20°C for 20 minutes, followed by hybridization.

### STARmap Library construction

For the STARmap-ISS method, 890 genes were selected based on a combination of highly variable genes and senescence related genes. STARmap-ISH genes were manually chosen from canonical and newly discovered senescence markers from snRNA-seq (**Table S1**). SNAIL probes were designed, and the probe library was constructed according to Wang et al.^8^. The probes were dissolved in ultrapure RNase-free water and pooled to a final concentration of 4 nM per probe. The probe mixture was heated to 40°C for 15 minutes and then equilibrated to 37°C. Tissue samples were removed from -20°C storage, equilibrated to room temperature, and treated with 10 mM Tris pH 7.5 for 10 minutes. The samples were washed with PBSTR (0.1% Tween-20, 0.1 U/μL SUPERase•In in PBS) and incubated in hybridization buffer (2X SSC, 10% formamide, 1% Tween-20, 20 mM ribonucleoside vanadyl complex, 0.1 mg/mL yeast tRNA, 0.1 U/μL SUPERase•In, SNAIL probes (4 nM per probe for STARmap-ISS, 10nM per probe for STARmap-ISH) at 40°C with gentle shaking for 48 hours. The samples were washed twice with PBSTR for 20 minutes at 37°C and then with 4X SSC in PBSTR for 20 minutes at 37°C, followed by a rinse with PBSTR at room temperature.

The SNAIL padlock probes annealed to the samples were ligated by incubating with a T4 DNA ligation mixture (1:10 dilution of T4 DNA ligase, 0.2 mg/mL BSA, 0.5 U/μL SUPERase•In) for 2 hours at room temperature with gentle agitation, followed by two 5-minute washes with PBSTR. The samples were then incubated in the RCA mixture (1:50 dilution of Phi29 DNA polymerase, 250 μM dNTP, 20 μM 5-(3-aminoallyl)-dUTP,

0.2 mg/mL BSA, 0.2 U/μL SUPERase•In) for 2 hours at 30°C with gentle agitation, followed by two 5-minute washes with PBSTR. Subsequently, the samples were treated with 25 mM acrylic acid NHS ester for 2 hours at room temperature with agitation, rinsed with PBST (0.1% Tween-20 in PBS) once, and incubated with monomer buffer (4% acrylamide, 0.2% bis-acrylamide, 2X SSC in H2O) for 15 minutes for polymerization pre-treatment. The buffer was removed, and 30 μL of monomer mixture (0.1% ammonium persulfate, 0.1% tetramethylethylenediamine in monomer buffer) was directly added to the center of each sample, which was immediately covered with a coverslip (coated with Gel-Slick Solution according to the manufacturer’s instructions) and allowed to polymerize at room temperature for 1 hour. The tissue-gel hybrid was washed with PBST twice and cleared by proteinase K digestion mixture (50 mM Tris pH 7.5, 100 mM NaCl, 1% SDS, 0.2 mg/mL proteinase K in H2O) at 37°C overnight. On the following day, the samples were treated with a dephosphorylation mixture (1:100 dilution of shrimp alkaline phosphatase, 0.2 mg/mL BSA, 1:10 dilution of CutSmart buffer in H2O) and rinsed with PBST.

### Imaging and Sequencing

For *in situ* hybridization detection, the 19-nt fluorescent oligo complementary to the DNA amplicon was diluted to 100 nM in 1X SSC dissolved in PBST, and samples were incubated at room temperature for 30 minutes, then washed with PBST three times for 5 minutes each before imaging. For *in situ* sequencing detection, the protocol was followed according to Wang et al.^8^. Each sequencing cycle started with treatment with stripping buffer (60% formamide, 0.1% Triton-X-100) for 5 minutes and triple-washing with PBST for 5 minutes. The samples were then incubated with a sequencing mixture (0.2 mg/mL BSA, 10 μM reading probe, 5 μM fluorescent decoding probe, 1:25 dilution of T4 DNA ligase) for 3 hours at room temperature. Subsequently, the samples were triple washed with washing and imaging buffers (2X SSC, 10% formamide) for 10 minutes before imaging. DAPI staining was performed after Cycle 6 of imaging for cell segmentation. Images were acquired using a Leica Stellaris 5 confocal microscope with a 405 nm diode, white light laser, and a 20x air objective (NA 0.75).

### Imaging preprocessing and registration

Images for individual tiles were deconvoluted using Huygens (23.04) to enhance image signals and suppress background noises. Next, we proceeded with image registration for different sequencing rounds, employing the 3D Fourier Transform implemented through the functions available in numpy.fft and Scipy. Cross-correlation between pairs of images at all translational offsets was computed, and the position associated with the highest correlation coefficient was identified. This position was then utilized to translate image volumes to compensate for the offset. During this procedure, the first sequencing round was used as reference, and subsequent rounds were registered to be aligned with it.

### Spot calling and decoding

After registration, individual dots were identified in each color channel on the first round of sequencing. Then, starfish (v0.2.2) was used to process sequencing signals. The registered images were converted into starfish format for downstream analysis.

Amplicon dots were identified by BlobDetector in starfish, where the signal of each dot was treated as a Gaussian kernel. BlobDetector convolved kernels of multiple sizes and picked the best fit for each spot. The minimum and maximum standard deviation of Gaussian kernels were set as 0.5 and 10.0 voxel volume, respectively, due to the predominant spot sizes falling within this specific range. The number of kernels was set as 10. Spots exhibiting local maxima in the lowest 10-20% of intensity values were eliminated to remove low-quality signals. Then, identified spots were decoded using PerRoundMaxChannel function to pick the channel with maximum signal intensity within voxel volume search radius for each round. A channel vector was generated for each spot by selecting channels from all six rounds. This vector was subsequently translated into a gene barcode using a barcode codebook, wherein each channel is linked with a 2-base sequence encoding. Spots with undetectable signals in certain rounds were excluded from the analysis, because spots with gene barcodes do not present in the barcode codebook.

### Cell Segmentation

Cell segmentation was performed on DAPI channel images using the pretrained 2D_versatile_fluo model from StarDist^105^. After segmentation, the centroids and areas of the identified cells were extracted. These centroids served as markers for further refinement using a watershed algorithm. The resulting segmented image was then used to assign decoded transcripts to individual cells. Transcripts that were not assigned to any cells were excluded from downstream analysis.

### Single-cell Data Analysis

The resulting count matrices were analyzed using Seurat (v4.3.0) in R (v4.2.2)^97^. For each sample, cells with fewer than 5 transcripts or 5 genes were filtered out. The remaining cells were normalized using the SCTransform function (with vst.flavor= “v2” among other default parameters), followed by PCA. Default parameters were used for the FindNeighbors, FindClusters, and RunUMAP functions. Subsequently, old and young mouse lung datasets were integrated separately. The functions SelectIntegrationFeatures (with nfeatures= 890 among other default parameters), PrepSCTIntegration, and FindIntegrationAnchors (with normalization.method= “SCT” among other default parameters) were used. Finally, the datasets were integrated using the IntegrateData function (with normalization.method= “SCT”).

### Label Transfer

STARmap cell-type annotation was performed by transferring labels from single-cell RNA-seq reference data using Seurat and validated the clusters using the collection of genes used for single-cell RNA-seq. The single-cell RNA-seq dataset was split into old and young samples and used to annotate the STARmap old and young cells, respectively. Initially, the single-cell RNA-seq datasets were normalized using the SCTransform function (with vst.flavor= “v2”, variable.features.n= 3000, min_cells= 1, return.only.var.genes= FALSE among other default parameters). The FindTransferAnchors function (with normalization.method= “SCT”, using all STARmap-detected genes as features, and other default parameters) was used to identify anchors between STARmap and single-cell RNA-seq objects. The TransferData and AddMetaData functions were then used to transfer and add cell-type annotations.

Finally, the MapQuery function was used to project the STARmap integrated object onto the single-cell RNA-seq UMAP embedding.

The identification of *p21^+^* senescent cells was conducted using Seurat subsetting on the expression level (*p21^+^*, *p16/p21^+^*>0)

### STARmap-ISS Imputation

The imputation of unmeasured genes was conducted by learning from single-cell RNA-seq data using Tangram. Tangram maps single-cell RNA-seq expression onto spatial transcriptomics data using a deep learning approach^71^. Both single-cell RNA-seq and STARmap datasets were normalized using the scanpy.pp.normalize function. The mapping between single-cell RNA-seq and STARmap data was performed using all genes decoded by STARmap as training genes. Finally, the imputation was completed with the project_genes function.

### Raman Imaging

A custom built NIR confocal Raman microscopy system was used for Raman data acquisition. The system was equipped with 785 nm wavelength Ti: Sapphire laser (3900S, Spectra Physics) and beam was filtered by a laser line filter (BPF, LL01 785 12.5, Semrock) and redirected to the dual axes galvanometer mirrors. High-speed XY scanning was performed by the galvanometer mirrors (CT 6210, Cambridge Technology) and large area scanning was performed by XY motorized stage (MS-2000, ASI Imaging). A 0.95 NA objective lens (Olympus UPLSAPO40X2 40X/0.95) was used to both focus the laser light onto the sample and to collect the back scattered light. A piezo actuator combined with a differential micrometer (DRV517, Thorlabs) was used to perform the coarse and fine adjustments, respectively, of the sample focus. A flip mirror was placed after the tube lens so that the sample focal plane from the incoherent transmission source can be observed using a video camera with 44 X magnification.

The backscattered Raman light from the sample passes through two dichroic mirrors (DM1: Semrock LPD01 785RU 25, DM2: Semrock LPD01 785RU 25×36×1.1) and was collected by a multi-mode fiber (Thorlabs M14L 01). The collected signal was delivered to the imaging spectrograph (Holospec f/1.8i, Kaiser Optical Systems) and detected by a thermoelectric cooled, back illuminated and deep depleted CCD (PIXIS: 100BR_eXcelon, Princeton Instruments). Data acquisition board (PCI 6251, National Instruments) and MATLAB 2022 software (Mathworks) were used to control the system, acquire the data, and analyze the data. Further details on how the confocal Raman measurements were performed are described previously^106^.

Tissue slices from mouse lung and skin are placed on top of quartz coverslips in the Petri dish with quartz coverslip bottom (SF-S-D12, WakenBtech). The dish is filled with 2mL of PBSR and wrapped by parafilm. The optical resolution limit of the system is 0.4 µm, and we used a sampling pixel size of 3 µm, this corresponds to an effective sampling-limited resolution of approximately 6 µm^107^. Raman field of view is guided by bright field imaging. For each sample, we selected a field of view in the range of approximately 750–1000 µm × 750–1000 µm, resulting in roughly 250–333 × 250–333 spectral measurements (with 3 µm pixel size).

### Spectrum Preprocessing

The obtained Raman imaging data measured the fingerprint region of Raman spectra (600-1800 cm⁻¹), capturing 873 spectral dimensions, where most of the signatures from key biomolecules such as proteins, nucleic acids, and metabolites are found. For the original Raman spectrum data, we implemented basic data processing steps using the Rampy python library, including cosmic-ray removal, smoothing by Savitzky-Golay filters, and background removal to reduce the artifact biological effects and noise deriving from the imaging process. To normalize the Raman imaging data, the area under the curve between the wavenumbers 1630 and 1700 cm⁻¹ was calculated using Simpson’s rule, since this region represents the amide I band, primarily indicative of protein secondary structures such as alpha-helices and beta-sheets. The intensity of each spectrum was then divided by this area, standardizing intensities for comparative analysis across samples. This spectrum normalization approach ensures that variations due to instrumental differences or sample concentration are minimized, allowing for more accurate cross-sample comparisons and robust biomolecular characterization.

### Raman-STARmap Registration

Due to inherent technological differences, Raman and STARmap tissue imaging techniques exhibit variations in resolution (Raman ∼ 2.5 µm, STARmap ∼ 0.284 µm) and fields of view (FoVs). In this study, Raman imaging covers approximately 20% of the area imaged by STARmap for each sample. Although STARmap is slower and costlier, it enables sequencing and high-resolution imaging of large areas. Conversely, Raman imaging is faster and more economical, albeit limited to scanning smaller areas with a relatively lower resolution. For alignment between Raman and STARmap images, we targeted the Raman peak at 791 cm⁻¹, commonly identified as the DNA signal, and aligned it with the DAPI images from STARmap. By using MATLAB’s Control Point Registration function, cpselect, we manually selected corresponding cellular keypoints across both imaging modalities. This selection tool then generated a 3×3 transformation matrix to adjust the STARmap images to align with the Raman regions. The manual alignment process utilized a least-squares method, employing a modified two-dimensional version of Horn’s (1987) algorithm to account for differences in translation, scale, rotation, and reflection. For each Raman-STARmap paired sample, hundreds of keypoints were manually selected, and the fitgeotrans function in MATLAB was used to transform the STARmap image to match the Raman region. The imshowpair function was employed iteratively after every 20 keypoints to ensure satisfactory alignment.

### Raman Cell Segmentation

After registration, both Raman and STARmap images are aligned within the same FoVs. Given the relatively lower resolution of Raman imaging, directly implementing cell segmentation on these images presents challenges. Consequently, we averaged the cell segmentation results from STARmap DAPI images to delineate the Raman signal within each STARmap cell boundary. Specifically, we used segmentation results previously obtained with StarDist for downstream analysis, aligning them with the Raman image using the same transformation matrix. This method yields an aligned cell segmentation that can be directly applied to segment the Raman image effectively.

Finally, we aggregated Raman spectra confined within the cell boundaries, obtaining an 873-dimensional Raman spectroscopic representation for each cell. The Raman spectral features are then paired with STARmap gene expression profiles and cell types at the single-cell level.

### Image and Peak Differential Analysis

For Raman image analysis, we compared cell sizes based on the number of Raman pixels within each cell boundary and retained only results with statistically significant differences, as indicated by an FDR-adjusted p≤ 0.05 using the Mann-Whitney U test^108^ We also calculated the Raman pixel intensity for each cell; the cell intensity was determined by averaging the peak intensities across all cells, adhering to the same statistical test and p threshold. Following a method analogous to the DEG analysis in snRNA-seq data, we identified significant spectral differences in the Raman data, which we refer to as Differentially Prominent Peaks (DPPs). For the analysis of DPPs, we employed a DEG-like approach to compare Raman peak intensities between senescent and non-senescent cells, aiming to detect peaks with varying intensities. We calculated FDR-adjusted p using the Wilcoxon rank-sum test and selected peaks with p≤ 0.05 as significantly different at the global level analysis and p≤ 0.1 at the celltype level due to the limited cell numbers. These peaks were then categorized into increased or decreased groups based on their log2FC values from normalized spectra. We selected the top 30 increased and decreased peaks based on the adjusted p and log2FC value from each group for further analysis. All Raman peak assignments were based on established annotations.

### Barcode Generation

We generated barcode plots to compare Raman intensities (and snRNA-seq gene expression) between senescent and non-senescent cells. First, the feature values were averaged for both cell groups and were normalized by dividing by the maximum value to ensure consistent scaling. Horizontal bars were then plotted for each feature, with gene features shown in red and Raman features in blue. The bars were positioned with a slight offset to prevent overlap, and representative features were labeled and annotated for their biological meanings. The bars are ranked by their average normalized Raman intensities or gene expression values, and y-axis labels indicated the cell groups compared. For integration analysis, the gene expression and Raman intensities were normalized separately, and we selected the top 60 multimodal features identified by the classifier from the combined DEG and DP sets using the same method to generate barcodes.

### Raman and snRNA-seq Integration

We integrated snRNA-seq and Raman imaging data using STARMap as the anchor. Specifically, we imputed the STARMap gene expression profiles to whole transcriptomic resolution for each cell using Tangram. These profiles were then paired with corresponding Raman peak features for each individual cell. Subsequently, we selected all the DEGs identified by snRNA-seq and the top 60 DEPs (top 30 increased and top 30 decreased, respectively) identified by Raman from the previous analysis and concatenated these features to create a multimodal differential feature combination for integration analysis.

### Machine Learning and Feature Importance Analysis

For both individual gene expression and Raman peak features, as well as their multimodal combinations, we built a Random Forest classifier to distinguish between senescent and non-senescent cells. Due to the significant class imbalance between the senescent and non-senescent cells, we downsampled the non-senescent class to equalize the number of cells in both categories. For different settings, we randomly shuffled the data and used z-score normalization to further standardize the multimodal features, splitting 70% for training and 30% for testing. The default parameters are used for Random Forest in scikit-learn, with n_estimators set to 500. After training, we reported the accuracy (ACC), area under the ROC curve (AUC), F1 score, Precision, and Recall on the test data. Since the inherent feature importance in Random Forest does not explicitly indicate the positive or negative influence related to the classes, we additionally used Shapley values to calculate feature importance. The TreeExplainer function is used to build the shap explainer since Random Forest is a tree-based model. The concept of Shapley value, from cooperative game theory, offers desirable properties. They indicate whether features positively or negatively impact the prediction, with their magnitude reflecting the strength of the effect. This analysis helps us better understand how the combinations of Raman and gene expression features characterize senescent cells. For correlation analysis, we simply use the clustermap function with hierarchical clustering in the Seaborn package.

## Extended Data

**Extended Data Fig.1.**
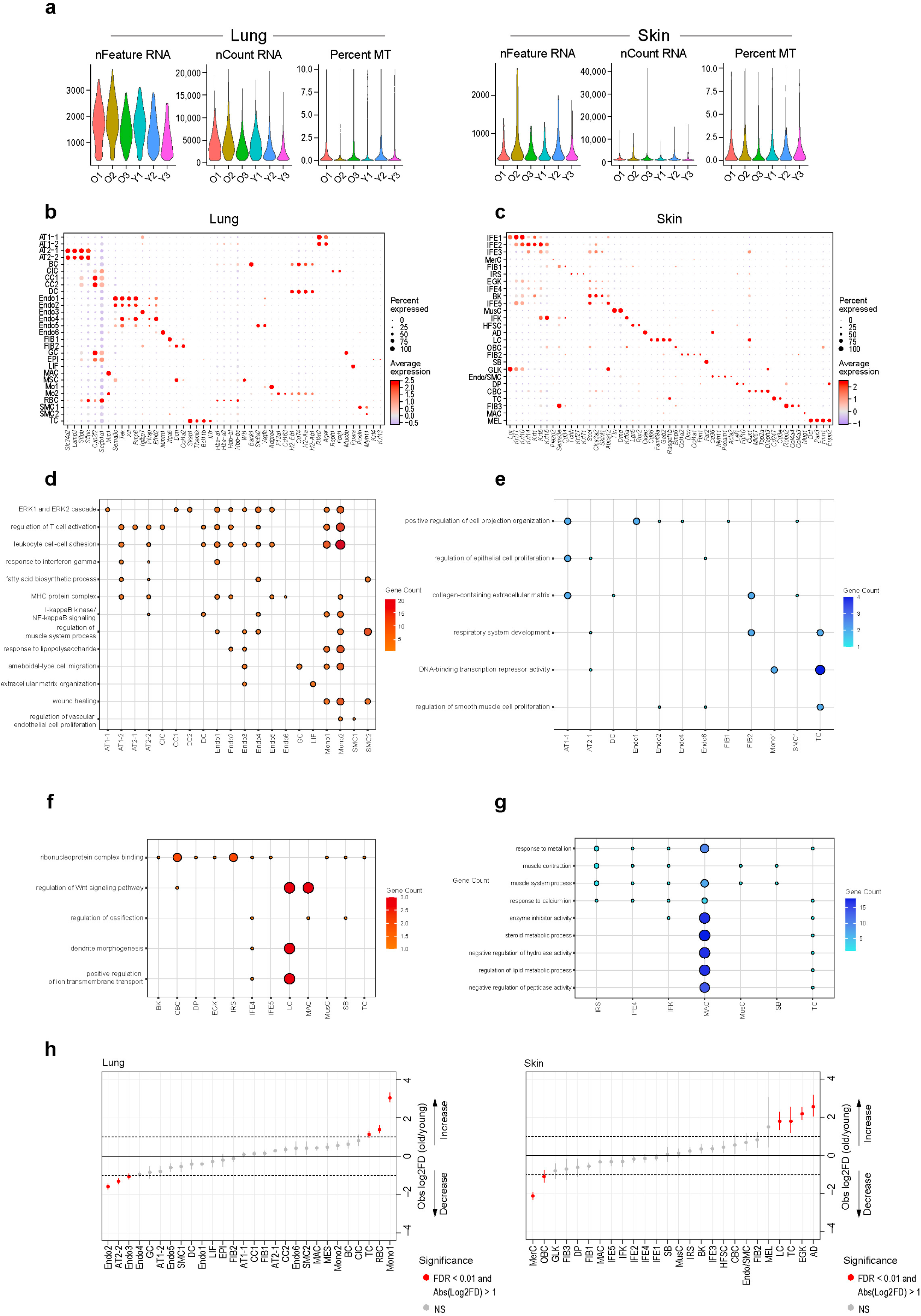
Further characterization of single nuclei transcriptomic data for aging analysis. **a**, snRNA-seq quality control metrics, separated by samples and tissues. Upper panel (lung), bottom panel (skin). Left: Number of genes detected per cell (nFeature_RNA) for each sample. Middle: Unique molecular identifiers per cell (nCount_RNA) for each sample. Right: Percentage of mitochondrial transcripts detected per cell (Percent.MT) for each sample. **b-c**, Dot plots representing the selected canonical marker genes expressed across main cell types in mouse lung (**b**) and skin (**c**). **d-e**, Representative Gene Ontology (GO) terms enriched among age-associated upregulated (**d**) and downregulated (**e**) DEGs across lung cell types. Bar length denotes the number of genes contributing to each term (“Gene count”). **f-g**, Dot plots showing representative GO terms enriched among age-associated upregulated (**f**) and downregulated (**g**) DEGs across skin cell types. Dot size indicates the number of genes contributing to each term (“Gene count”). **h**, Dot plots showing cell-type abundance changes in old mouse lung and skin relative to young tissues. Red dots with error bar indicate significant changes (FDR<0.01 and log2 fold change (Log2FC)>1), grey dots with error bar indicate no significant changes.

**Extended Data Fig.2.**
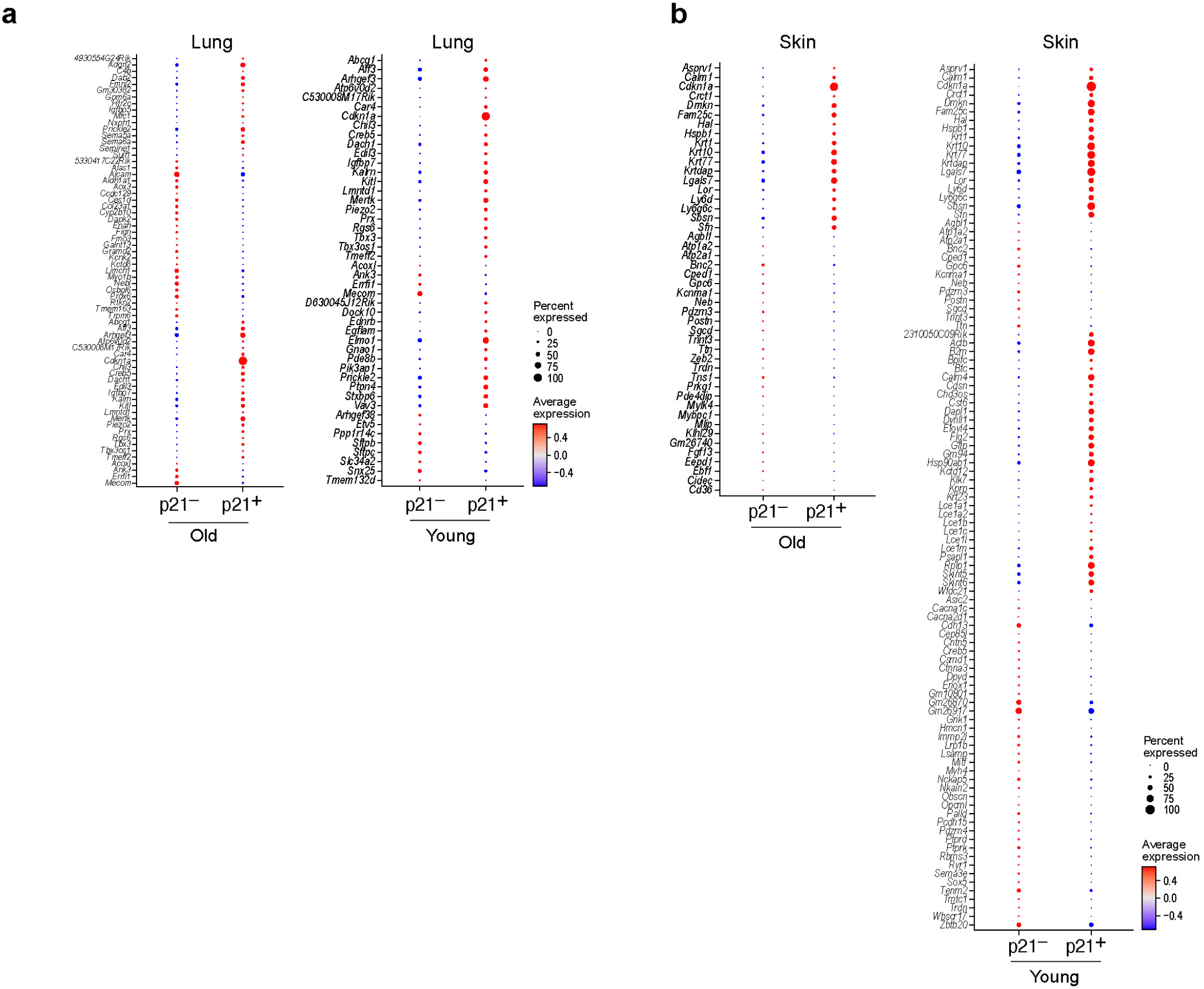
Further characterization of single nuclei transcriptomic data for senescence analysis. **a-b**, Dot plots showing DEGs specifically enriched in old or young samples in mouse lung (**a**) and skin (**b**). Average expression scores were calculated using log-normalized and scaled data (FDR<0.05). The numerical details are reported in Supplementary Table 6.

**Extended Data Fig.3.**
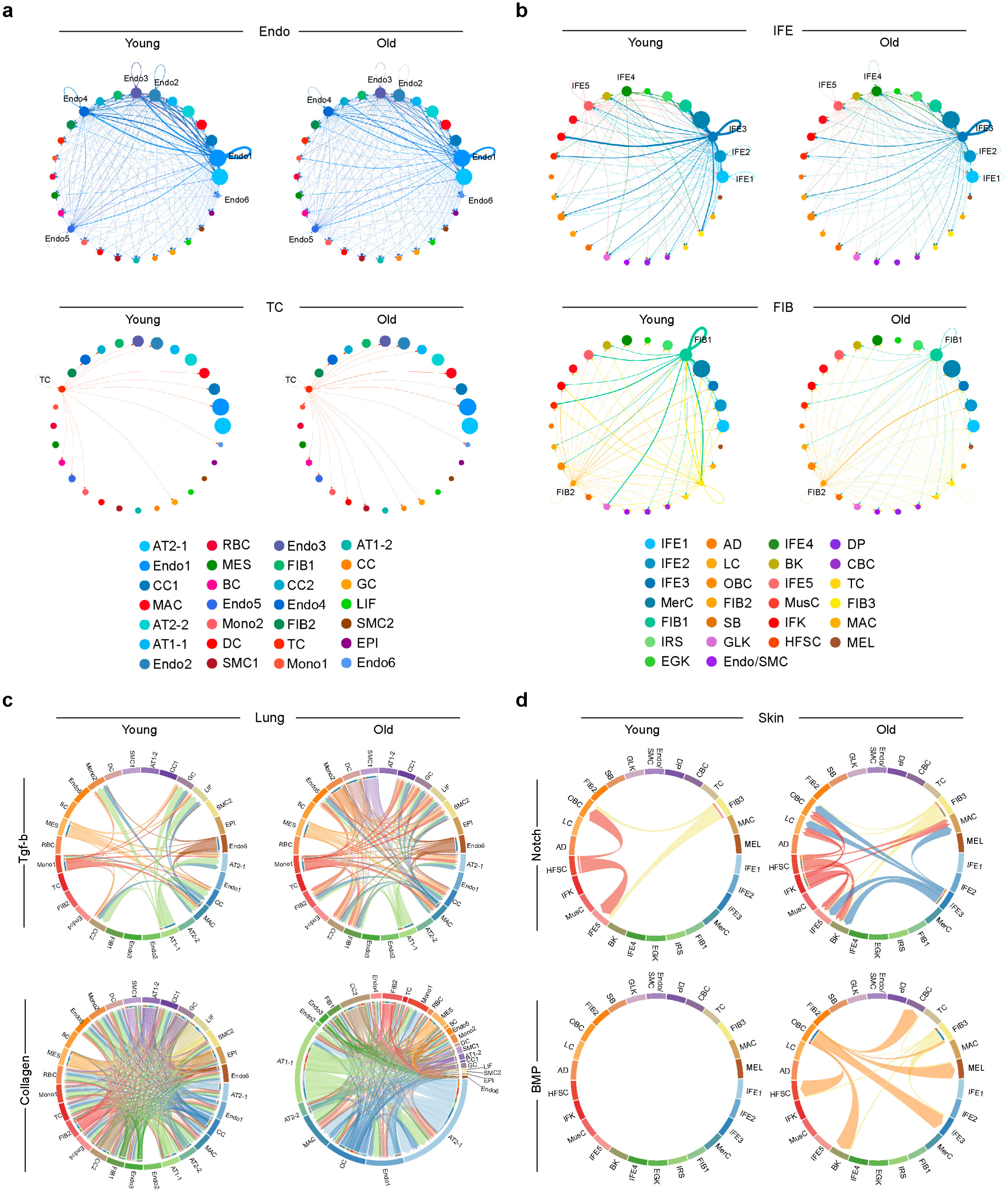
Cell type-resolved remodeling of intercellular communication with age in mouse lung and skin. **a**, Interaction net count plots showing cell-cell interactions between endothelial cell subtypes (top) or T cells (bottom) and other cell types. Line width reflects interaction strength, with thicker lines indicating a greater number of inferred ligand-receptor interactions between cell types. **b**, Interaction net count plots showing cell-cell interactions between interfollicular epidermis cell subtypes (top) or fibroblast cells (bottom) and other cell types. Line width reflects interaction strength, with thicker lines indicating a greater number of inferred ligand-receptor interactions between cell types. **c-d**, Chord plots showing the communication strength between different cell types through specific signaling pathways in mouse lung (**c**) and skin (**d**) at different ages. Arrow stem (signaling sender), arrowhead (signaling receiver), arrow width (signaling strength).

**Extended Data Fig.4.**
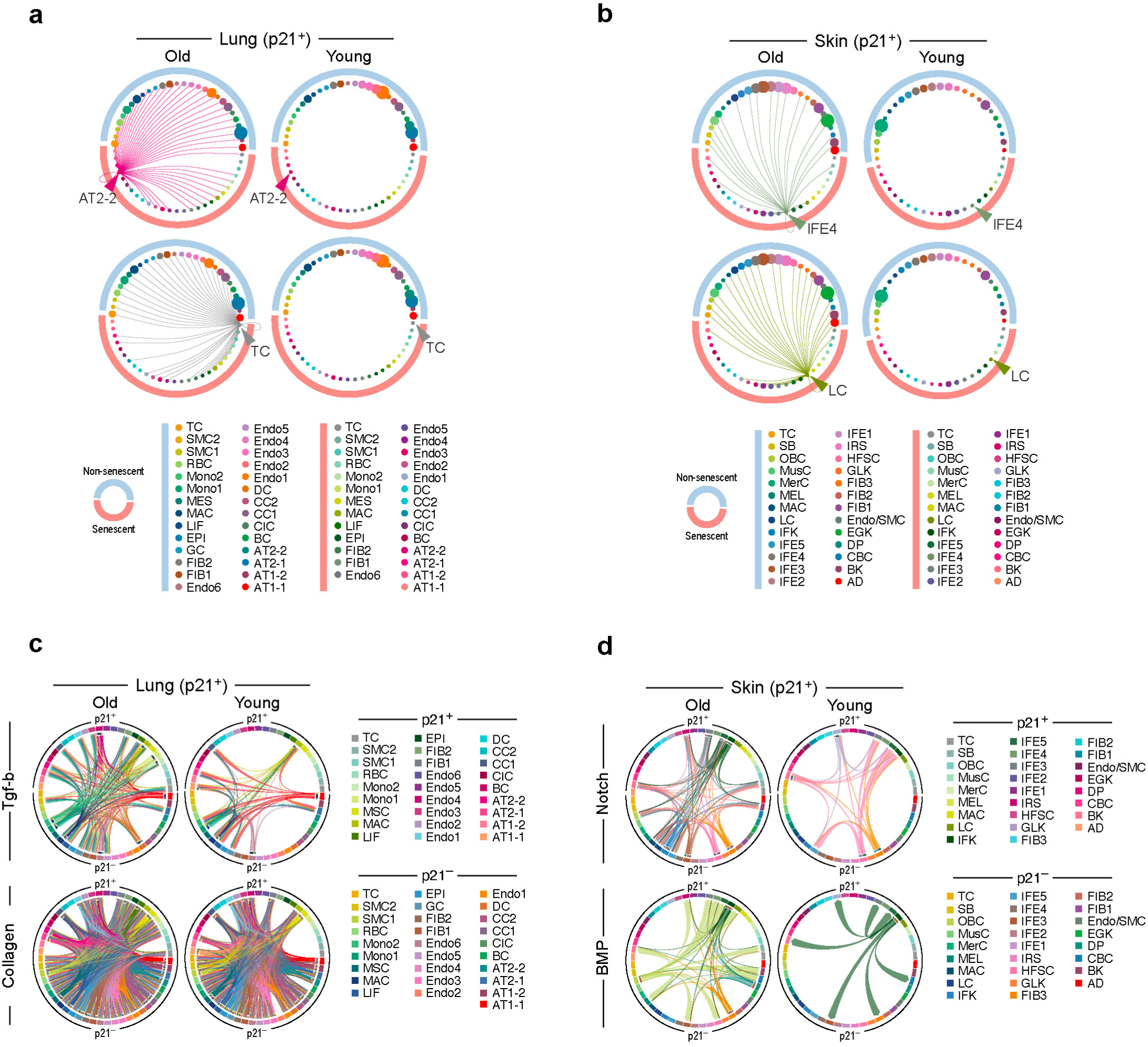
Cell type-resolved remodeling of intercellular communication between senescent and non-senescent cells in mouse lung and skin. **a-b**, Circle plots depicting the interactions of senescent AT2 and senescent T cells with other cell types in mouse lung (**a**), senescent IFE4 and senescent LC with other cell types in mouse skin (**b**) at different ages. Line width indicates the interaction strength. **c-d**, Chord plots showing the communication strength between different cell types through specific signaling pathways in mouse lung (**c**) and skin (**d**) at different ages. Arrow stem (signaling sender), arrowhead (signaling receiver), arrow width (signaling strength).

**Extended Data Fig.5.**
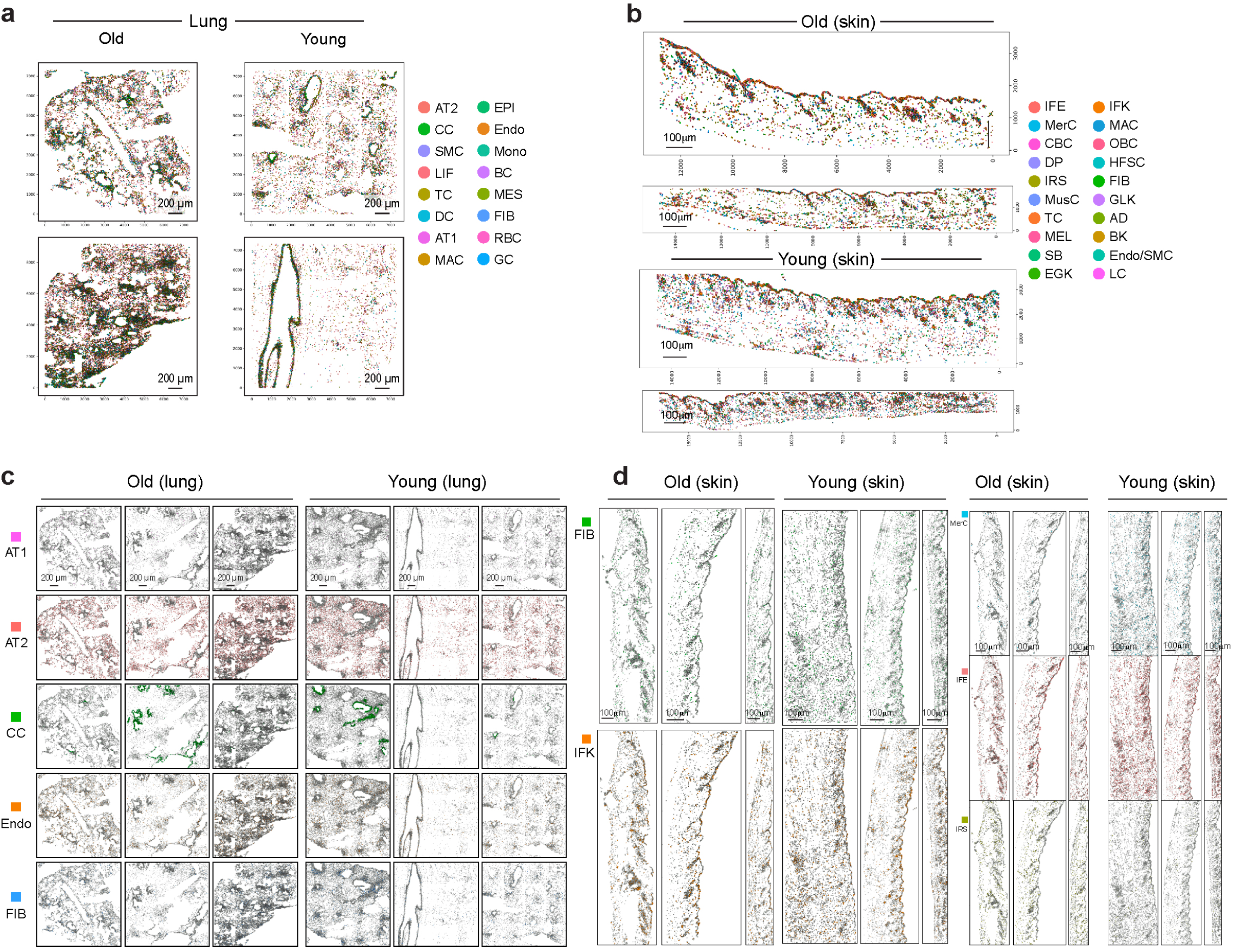
Spatial cell-type distributions revealed by STARmap-ISS in mouse lung and skin. **a-b**, snRNA-seq derived major cell types identified by STARmap-ISS (890 genes) via label transferring on old and young samples of mouse lung (**a**) and mouse skin (**b**). Cells are colored according to the cell type identification. Clusters composition can be found in Supplementary Table 2. **c-d**, Spatial localization of STARmap-ISS identified the top five most abundant cell types in mouse lung (**c**) and mouse skin (**d**).

**Extended Data Fig.6.**
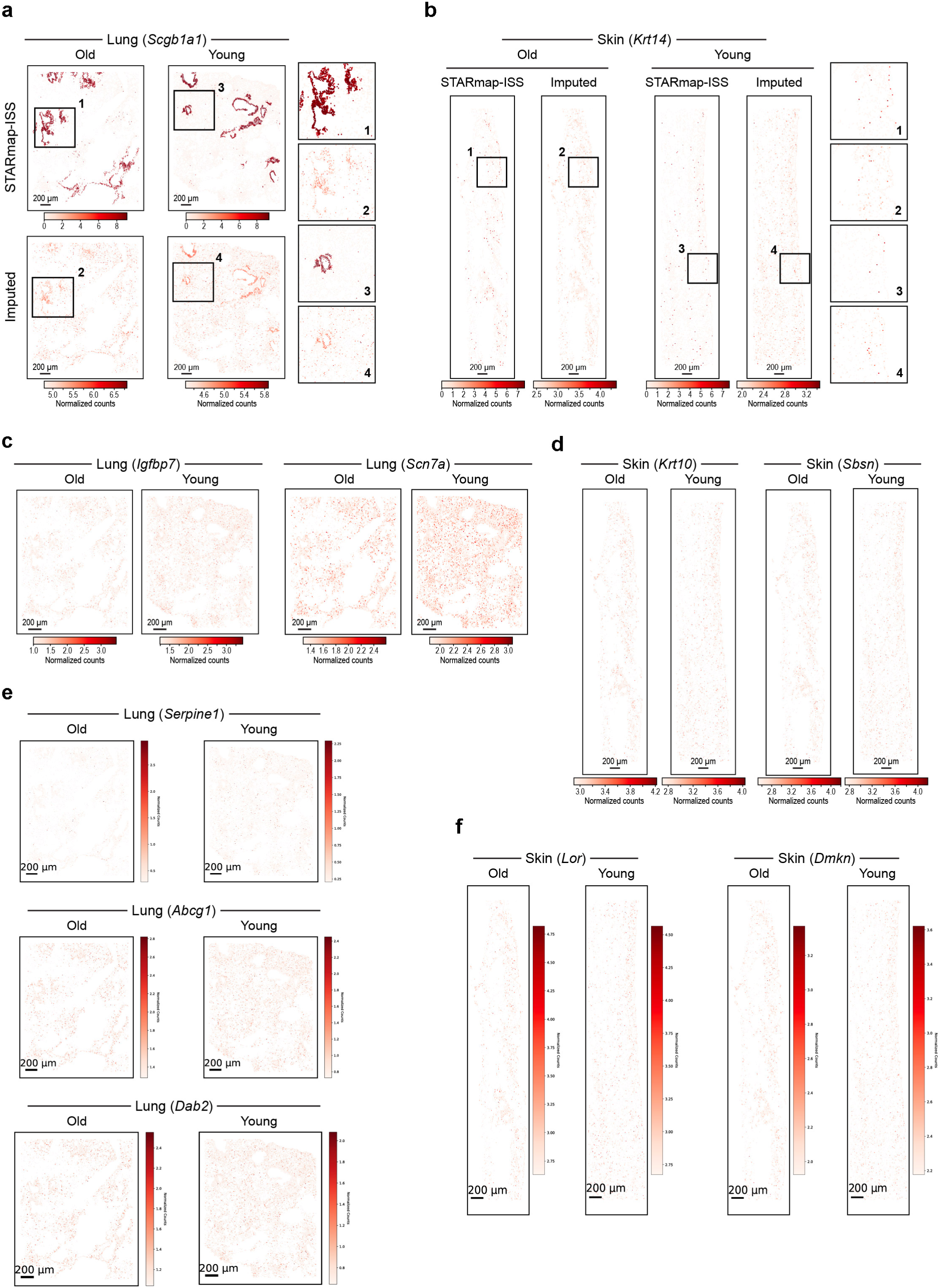
Spatial distribution of imputed senescence and aging markers in mouse lung and skin. **a-b**, Comparison of STARmap-ISS and imputation-based gene expression (Imputed) for genes expressing in club cells (*Scgb1a1*) (**a**) and basal keratinocyte (*Krt14*) (**b**) across tissues and ages. Black squares indicated area 1, 2, 3, 4 were enlarged for better visualization on the right side of each panel. **c-f**, Spatial visualization of imputed gene expression in mouse lung (**c**, **e**) and mouse skin (**d**, **f**) at different ages.

**Extended Data Fig.7.**
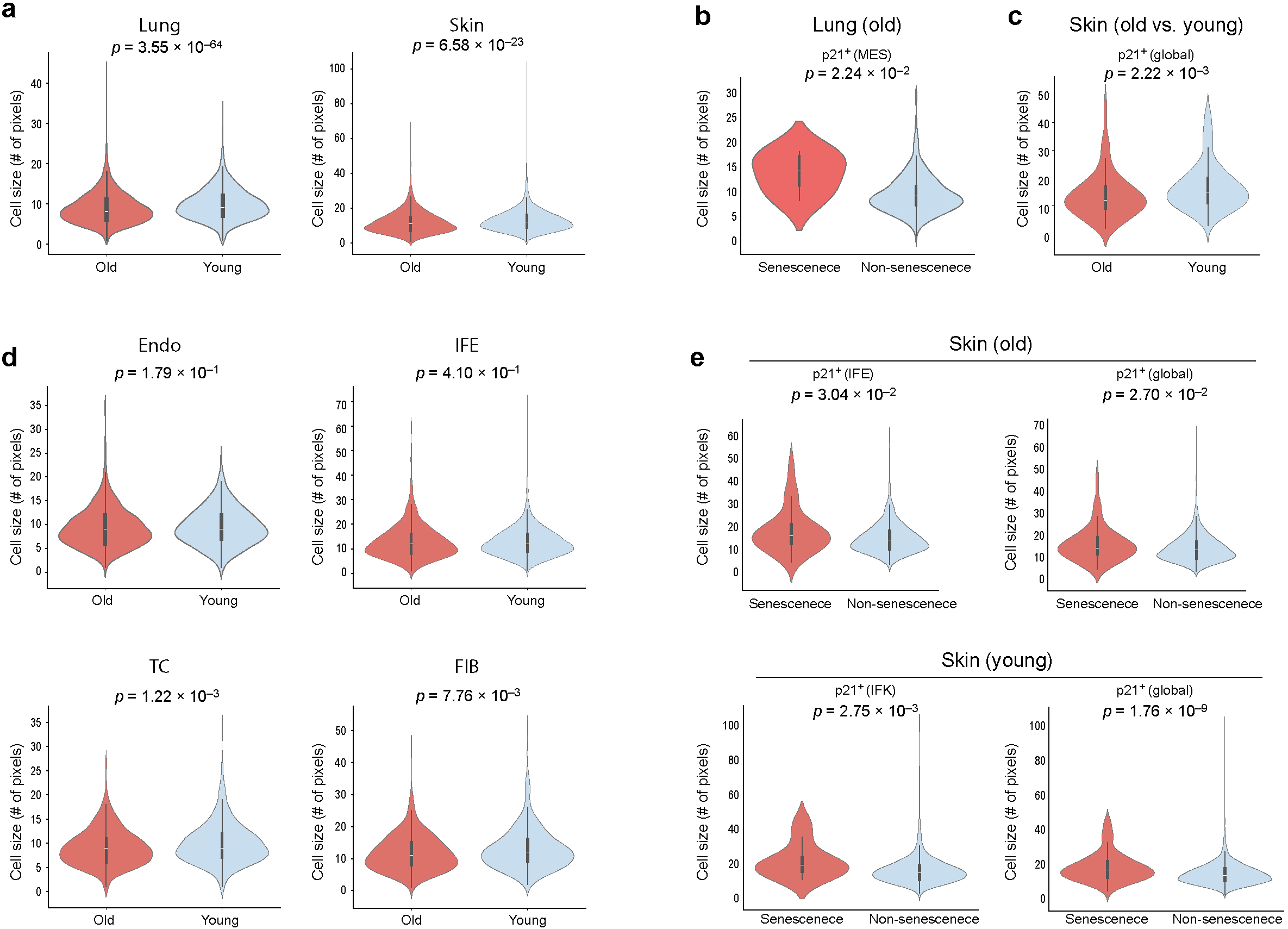
Comparative analysis of cell size differences across tissues and ages. **a**, Violin plots showing cell size quantified by the number of Raman pixels per cell in young and old mouse lung and skin at tissue level. **b**, Violin plot showing cell size measured by the number of Raman pixels in *p21*^+^ and *p21*^-^ mesenchymal cells from old mouse lung. **c**, Violin plots showing cell size measured by the number of Raman pixels, from young and old skin tissues in global *p21*^+^ cells. **d**, Violin plots showing cell size quantified by the number of Raman pixels per cell in young and old mouse lung and skin across different cell types. **e**, Violin plots showing cell size measured by the number of Raman pixels in *p21*^+^ labeled senescent cells and *p21*^-^ labeled non-senescent cells from young and old samples, analyzed global, IFE and IFK cells as indicated. p is calculated by the Mann–Whitney U test.

**Extended Data Fig.8.**
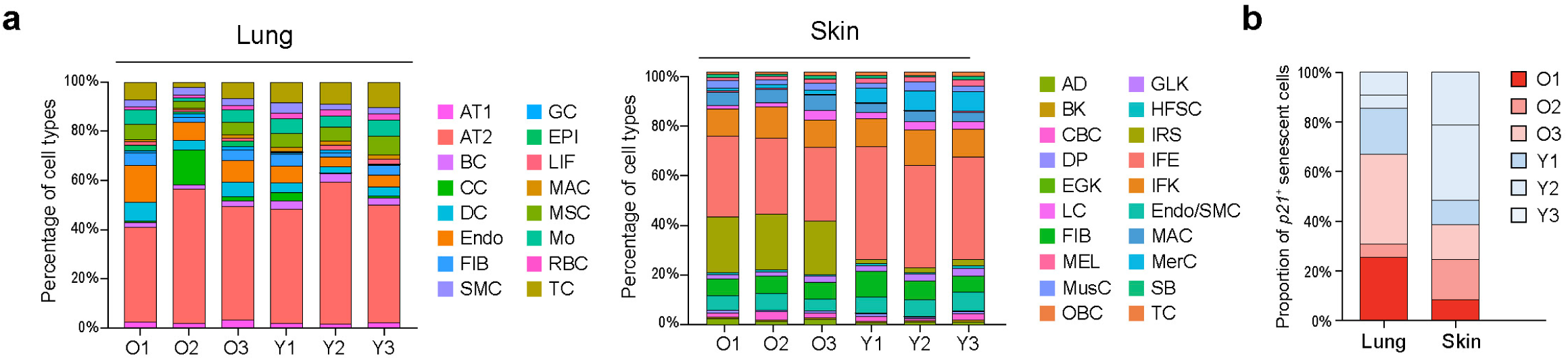
Cell-type composition and *p21*⁺ cell proportion in young and old mouse tissues. **a**, Bar graphs showing the percentages of all cell types identified from Raman imaging in mouse lung (left) and mouse skin (right). Cell types are colored according to STARmap-ISS cell-type clusters. **b**, Bar graph showing the proportion of *p21*^+^ senescent cells in mouse lung and skin across different age groups.

**Extended Data Fig.9.**
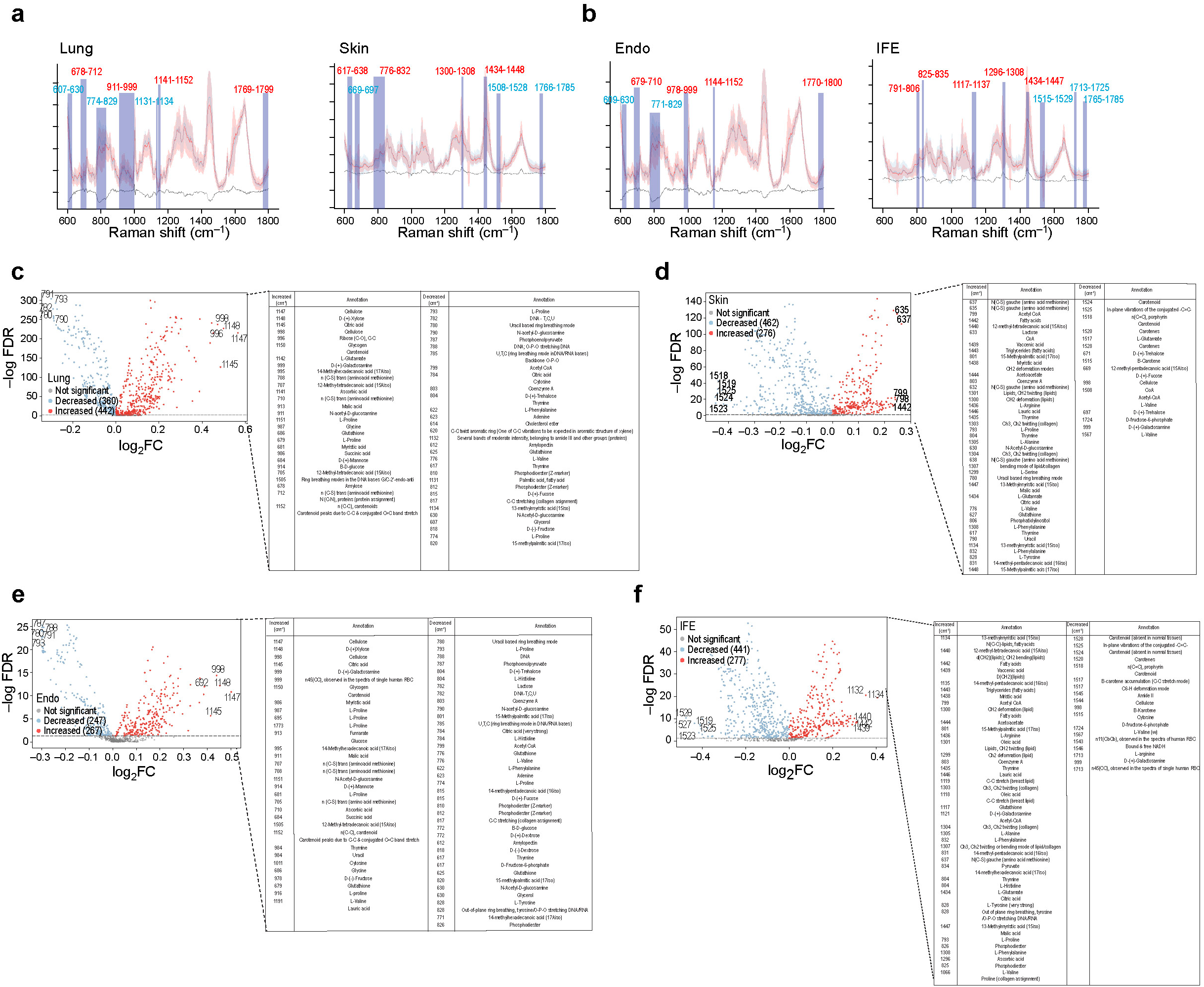
Raman spectral profiling of old and young cells across tissues at single-cell resolution. **a-b**, Comparison of the normalized Raman spectra (600-1800 cm⁻¹) of old cells (red line) and young cells (blue line) in mouse lung (**a**, left), mouse skin (**a**, right), Endo (**b**, left), and IFE cells (**b**, right). Dashed lines represent the differential spectrum, indicating variations between old and young cells. Shaded regions with Raman shift values indicate areas with significant DPs. Red numbers denote clusters enriched for increased DPs; blue numbers denote clusters enriched for decreased DPs. **c-f**, Volcano plots showing increased and decreased DPs, with corresponding biological annotations, in old versus young cells from mouse lung (**c**), lung endothelium (**e**), mouse skin (**d**), and IFE (**f**).

**Extended Data Fig.10.**
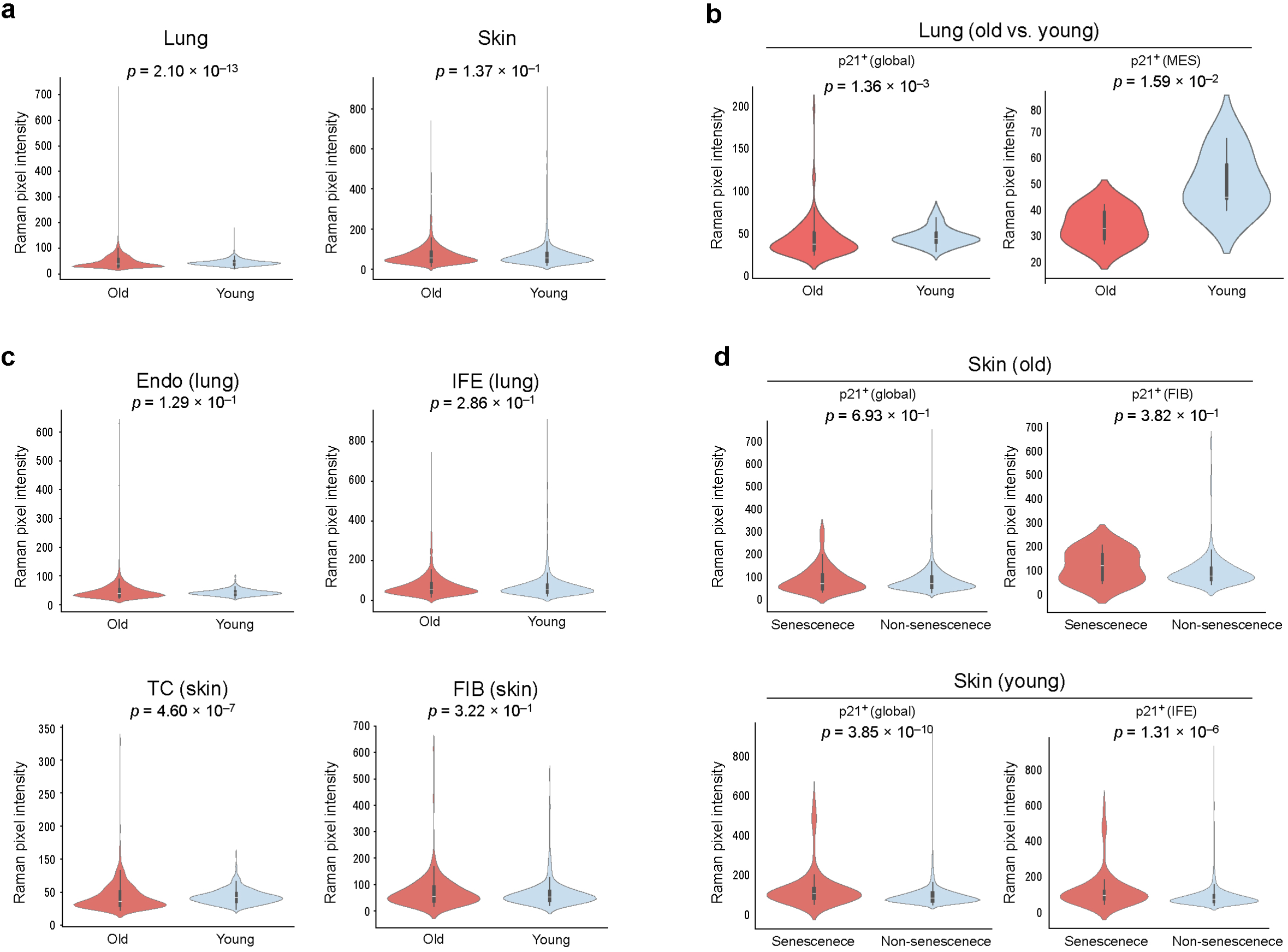
Comparative analysis of average Raman intensity differences across tissues and ages. **a**, Violin plots showing averaged Raman pixel intensity per cell in young and old mouse lung and skin at tissue level. **b**, Violin plots showing averaged Raman pixel intensity of all *p21*^+^ cells and *p21*^+^ mesenchymal cells from old and young mouse lung. **c**, Violin plots showing averaged Raman pixel intensity of young and old cells in different cell types across tissues. **d**, Violin plots showing averaged Raman pixel intensity of *p21*^+^ cells and *p21*^+^ cells at tissue or cell-type level in old and young skin. p is calculated by the Mann–Whitney U test.

**Extended Data Fig.11.**
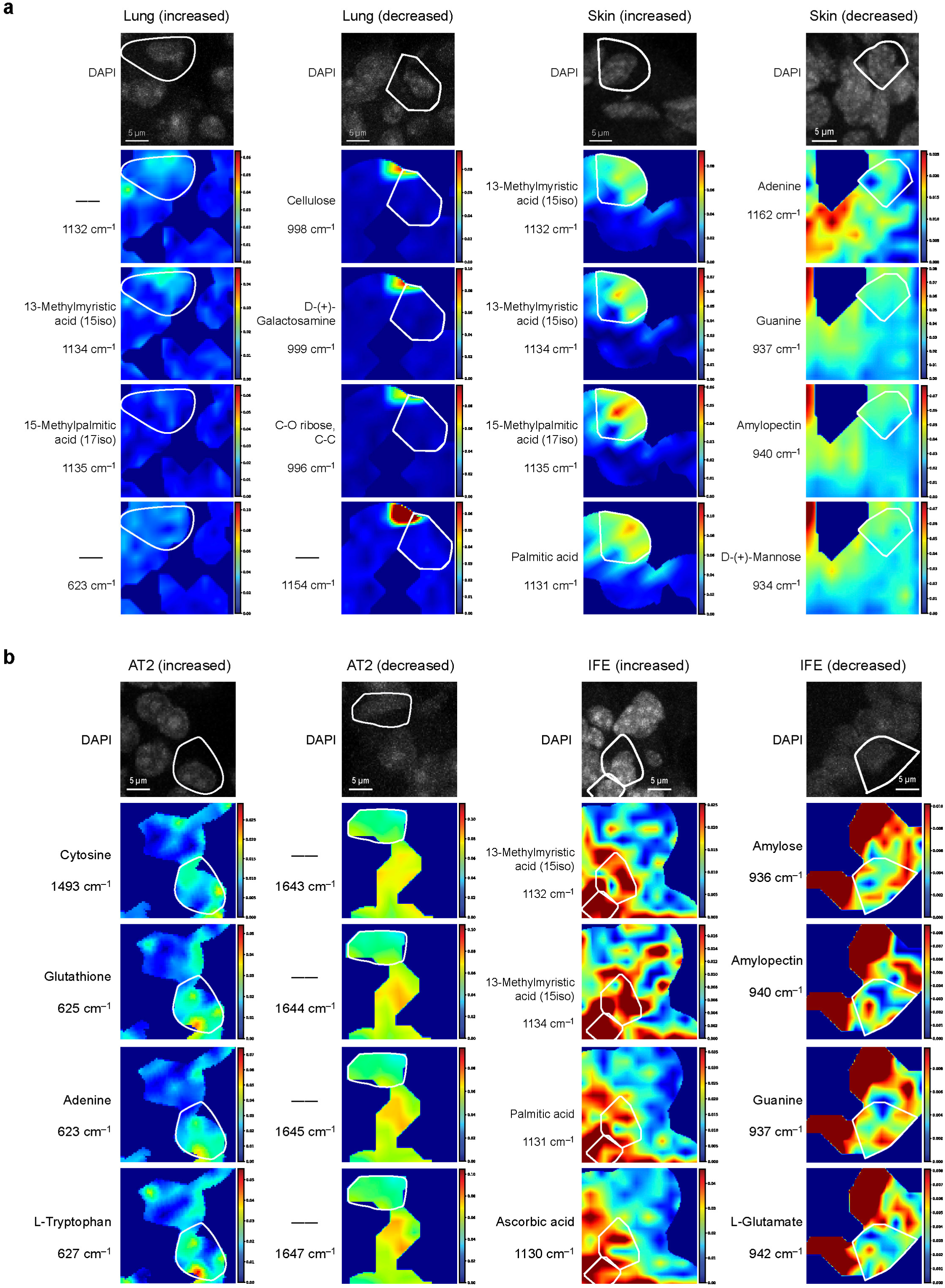
Visualization of differential Raman peaks at tissue and cell-type level. **a**, Spectral heatmaps showing Raman intensities of DPs alongside DAPI images from STARmap-ISS at the tissue level in mouse lung and skin, visualized at subcellular resolution. White outlines delineate the boundaries of senescent cells. **b**, Spectral heatmaps showing Raman intensities of DPs alongside DAPI images from STARmap-ISS at the cell-type level in AT2 cells (lung) and IFE cells (skin), visualized at subcellular resolution. White outlines delineate the boundaries of senescent cells.

**Extended Data Fig.12.**
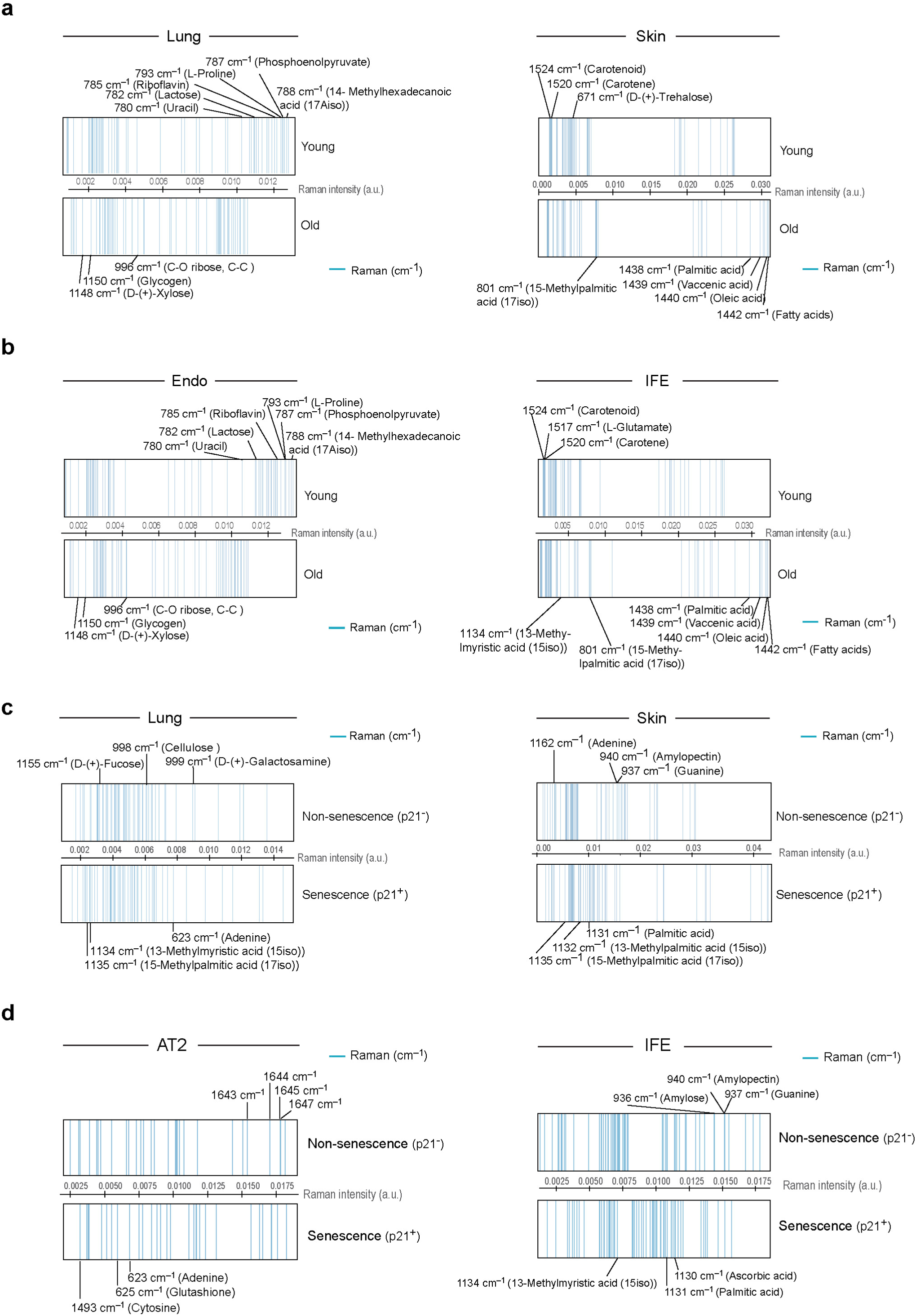
Raman spectral barcodes of aging and senescence across tissues and cell types. **a-b**, Stripe barcodes representing the most significant DPs in cells from old sample versus cells from young sample in mouse lung (**a**, left), mouse skin (**a**, right), Endo in mouse lung (**b**, left) and IFE in mouse skin (**b**, right). The barcodes display the intensity distribution of top 30 increased and top 30 decreased DPs, arranged based on the average peak intensities with representative DPs annotated for their biological meanings. The x-axis represents Raman intensity values. **c-d**, Stripe barcodes representing the most significant DPs in *p21*^+^ senescent cells versus *p21*^-^ non-senescent cells in mouse lung (**c**, left), mouse skin (**c**, right), AT2 in mouse lung (**d**, left) and IFE in mouse skin (**d**, right). The barcodes display the intensity distribution of top 30 increased and top 30 decreased DPs, arranged based on the average peak intensities with representative DPs annotated for their biological meanings. The x-axis represents Raman intensity values.

**Extended Data Fig.13.**
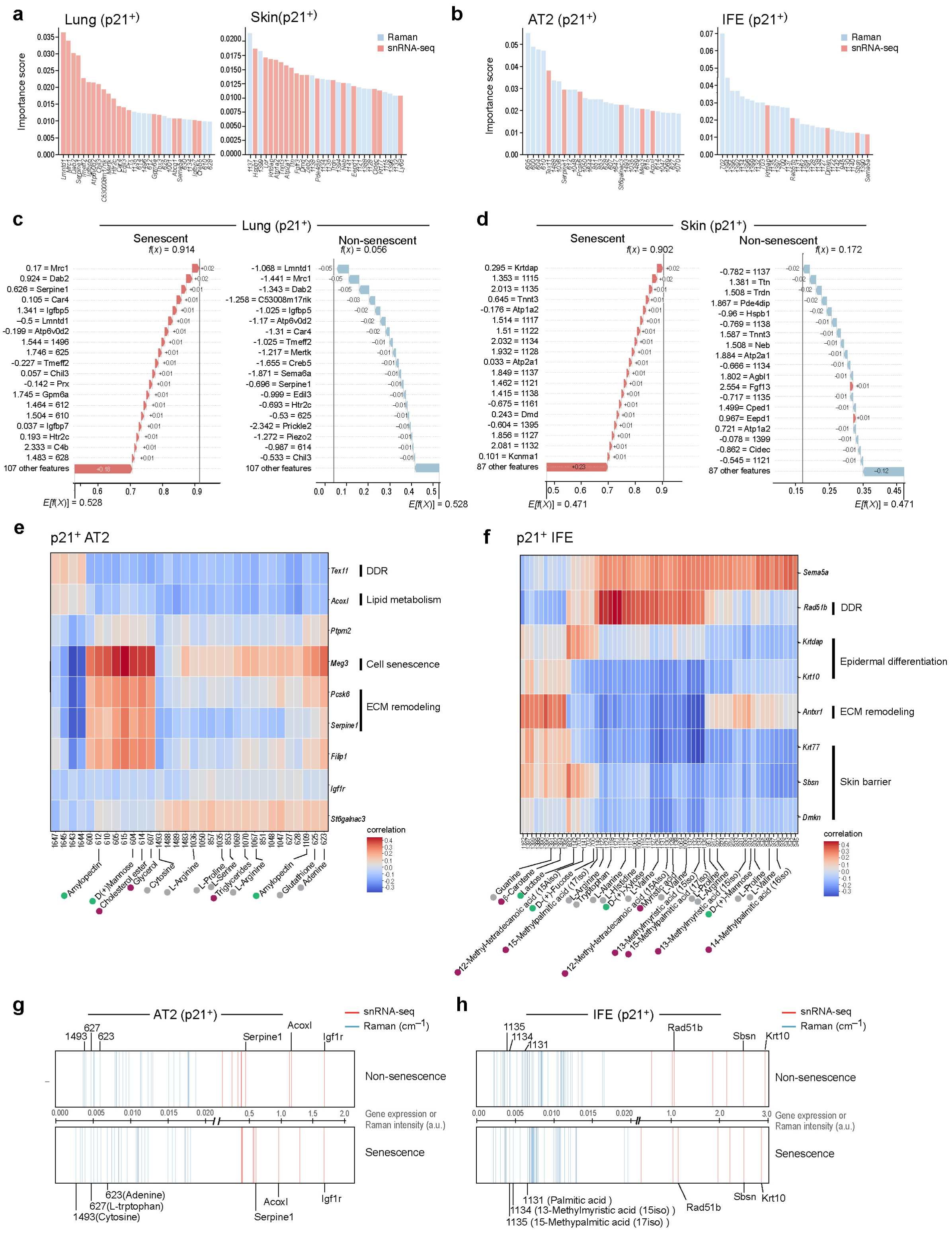
Further analysis of multimodal integration at cell-type level. **a**, Bar graphs showing the top 30 important features from integrated multimodal datasets using random forest in mouse lung (left) and mouse skin (right), ranked by their feature importances. Red, genes; Blue, Raman peaks. **b**, Bar graphs showing the top 30 important features from integrated multimodal datasets using SHAP method in mouse lung AT2 cells (left) and mouse skin IFE cells (right), ranked by their average SHAP values. Red, genes; Blue, Raman peaks. **c-d**, Waterfall graphs displaying the feature contributions of a randomly picked single cell to the prediction of senescent cells or non-senescent cells of the top ranked important features in mouse lung (**c**) and mouse skin (**d**). Red arrows with positive values indicate the predicting contribution to senescent cells, blue arrows with negative values indicate the predicting contribution to non-senescent cells. Numbers along with genes or peaks indicate the normalized Raman intensities or gene expression values of each feature. **e-f**, Heatmap showing the correlations between DPs (32 for AT2 cells, 60 for IFE cells) and DEGs (14 for AT2 cells, 8 for IFE cells) in *p21*^+^ senescent cells from AT2 cells (**e**) and IFE cells (**f**). Raman peaks and gene functions were manually annotated. **g-h**, Stripe barcodes representing the most significant DPs and DEGs in *p21^+^* senescent cells versus *p21^-^* non-senescent cells in mouse AT2 cells (**g**) and IFE cells (**h**). The barcodes display the top 32 DPs and top 14 DEGs for AT2 cells in mouse lung, and the top 60 DPs and top 8 DEGs for IFE cells in mouse skin. Raman peaks arranged based on average peak intensities with representative DPs annotated for their biological meanings. Genes arranged based on average gene expression level with representative genes highlighted.

**Extended Data Fig.14.**
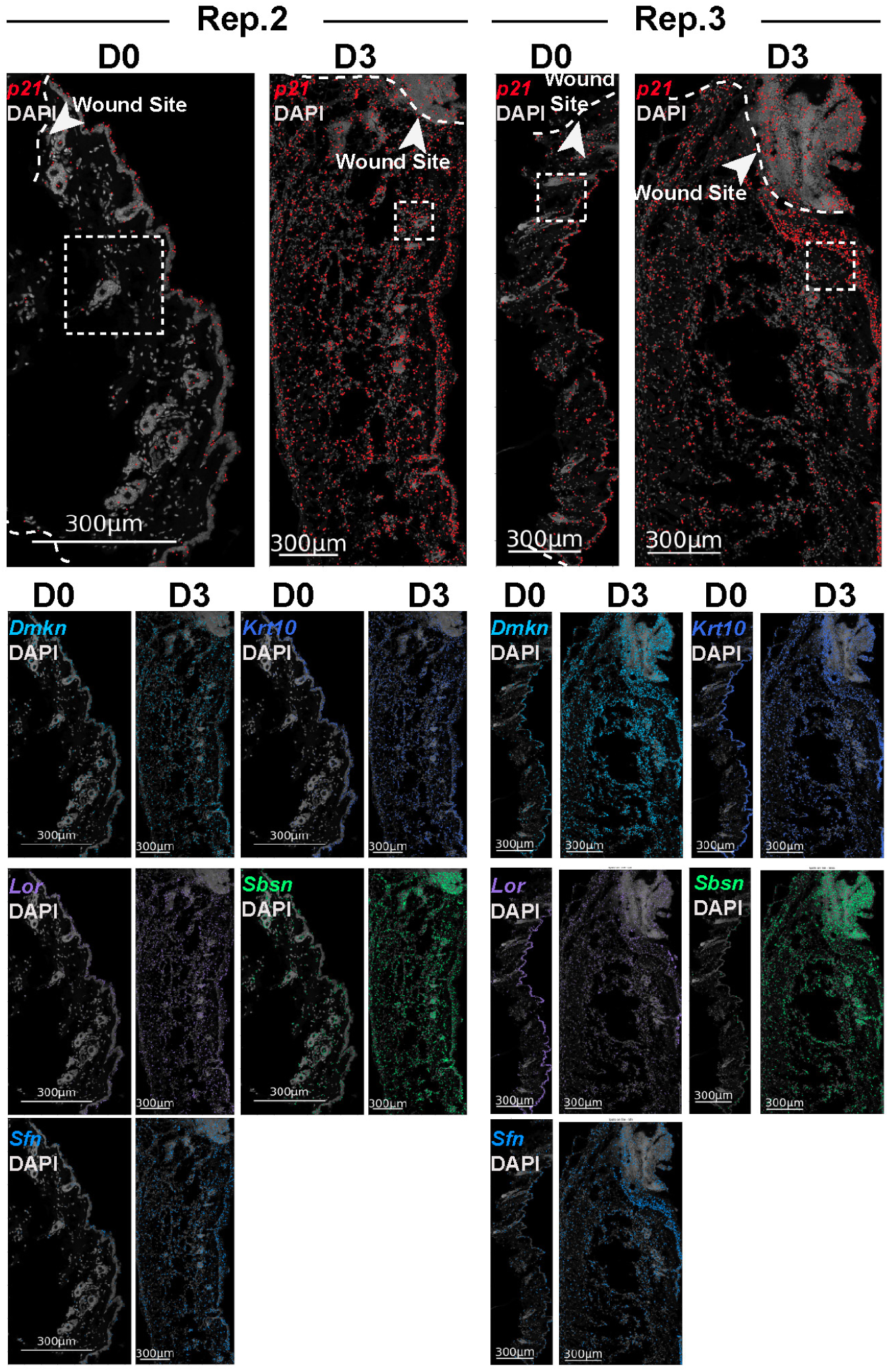
Spatial distribution of new senescence markers in wound model. Spatial distribution of newly identified senescence markers alongside the canonical marker *p21* at the wound site in old mouse skin at D0 and D3 post-injury.

## SUPPLEMENTAL INFORMATION

Table S1. STARmap-ISS and STARmap-ISH sequences used in this study

Table S2. Total cell number and senescent cell number from snRNA-seq, STARmap-ISS and integration analysis

Table S3. Raman pixel and average Raman intensity of different samples

Table S4. Differential Raman peaks of *p21^+^* vs. *p21^-^* cells and young vs. old cells in the lung, skin, AT2 and IFE

Table S5. The numerical details of metrics

Table S6. All cell-type makers and differentially expressed genes from snRNA-seq

